# Neural circuit mechanisms underlying context-specific halting in *Drosophila*

**DOI:** 10.1101/2023.09.25.559438

**Authors:** Neha Sapkal, Nino Mancini, Divya Sthanu Kumar, Nico Spiller, Kazuma Murakami, Gianna Vitelli, Benjamin Bargeron, Kate Maier, Katharina Eichler, Gregory S.X.E. Jefferis, Philip K. Shiu, Gabriella R. Sterne, Salil S. Bidaye

## Abstract

Walking is a complex motor program involving coordinated and distributed activity across the brain and the spinal cord. Halting appropriately at the correct time is a critical but often overlooked component of walking control. While recent studies have delineated specific genetically defined neuronal populations in the mouse brainstem that drive different types of halting^1–3^, the underlying neural circuit mechanisms responsible for overruling the competing walking-state neural activity to generate context-appropriate halting, remain unclear. Here, we elucidate two fundamental mechanisms by which *Drosophila* implement context-appropriate halting. The first mechanism (“walk-OFF” mechanism) relies on GABAergic neurons that inhibit specific descending walking commands in the brain, while the second mechanism (“brake” mechanism) relies on excitatory cholinergic neurons in the nerve-cord that lead to an active arrest of stepping movements. Using connectome-informed models^4–6^ and functional studies, we show that two neuronal types that deploy the “walk-OFF” mechanism inhibit distinct populations of walking-promotion neurons, leading to differential halting of forward-walking or steering. The “brake” neurons on the other hand, override all walking commands by simultaneously inhibiting descending walking promoting pathways and increasing the resistance at the leg-joints leading to an arrest of leg movements in the stance phase of walking. We characterized two ethologically relevant behavioral contexts in which the distinct halting mechanisms were used by the animal in a mutually exclusive manner: the “walk-OFF” pathway was engaged for halting during feeding, and the “brake” pathway was engaged for halting during grooming. Furthermore, this knowledge of the neural targets and mechanisms for halting, allowed us to use connectomics to predict novel halting pathways that could be relevant in other behavioral contexts.

## Main

Walking recruits distributed neural activity across the brain^7–12^ and spinal^13–16/nerve^ cord^17^. When an animal halts, the net output of this widely distributed neural activity dramatically changes to drive the arrest of leg stepping movements. Moreover, depending on the behavioral context, animals halt in different context appropriate ways. While a few specific neuronal types driving halting of walking have been recently described^1,2,18–21^, how walking related neural activity changes to manifest halting in a manner that is both biomechanically stable and appropriate to the behavioral context is currently unclear.

### Identification of specific neuronal types driving different types of halting

To identify neurons involved in execution of halting in *Drosophila*, we used red-shifted channelrhodopsin, CsChrimson^22^, to optogenetically activate specific neuronal types innervating the sub-esophageal zone (SEZ^23^) of the fly brain that houses the neurons close to and including the descending output of the brain^24^ (considered to be analogous to mammalian brainstem^25^). Through this neural activation screen (extended version of a previously published screen^26^, see Methods) we identified a set of 11 genetic drivers that were both relatively sparse and caused halting without gross motor deficits like paralysis or uncoordinated steps (Extended Data Fig. 1a). Among these lines, using intersectional genetics, we could unambiguously pinpoint three causal neuronal types for halting (Methods and Extended Data Fig. 1). We identified or generated split-Gal4 reagents that targeted these cell types with little to no ectopic expression and drove robust halting (Fig. 1a-c). Using image database search tools^27,28^ and coordinate transformation tools^29^, we were able to successfully identify these neurons in the brain^5^ and nerve-cord^30–32^ connectomes (Fig. 1e-f).

**Fig. 1:**
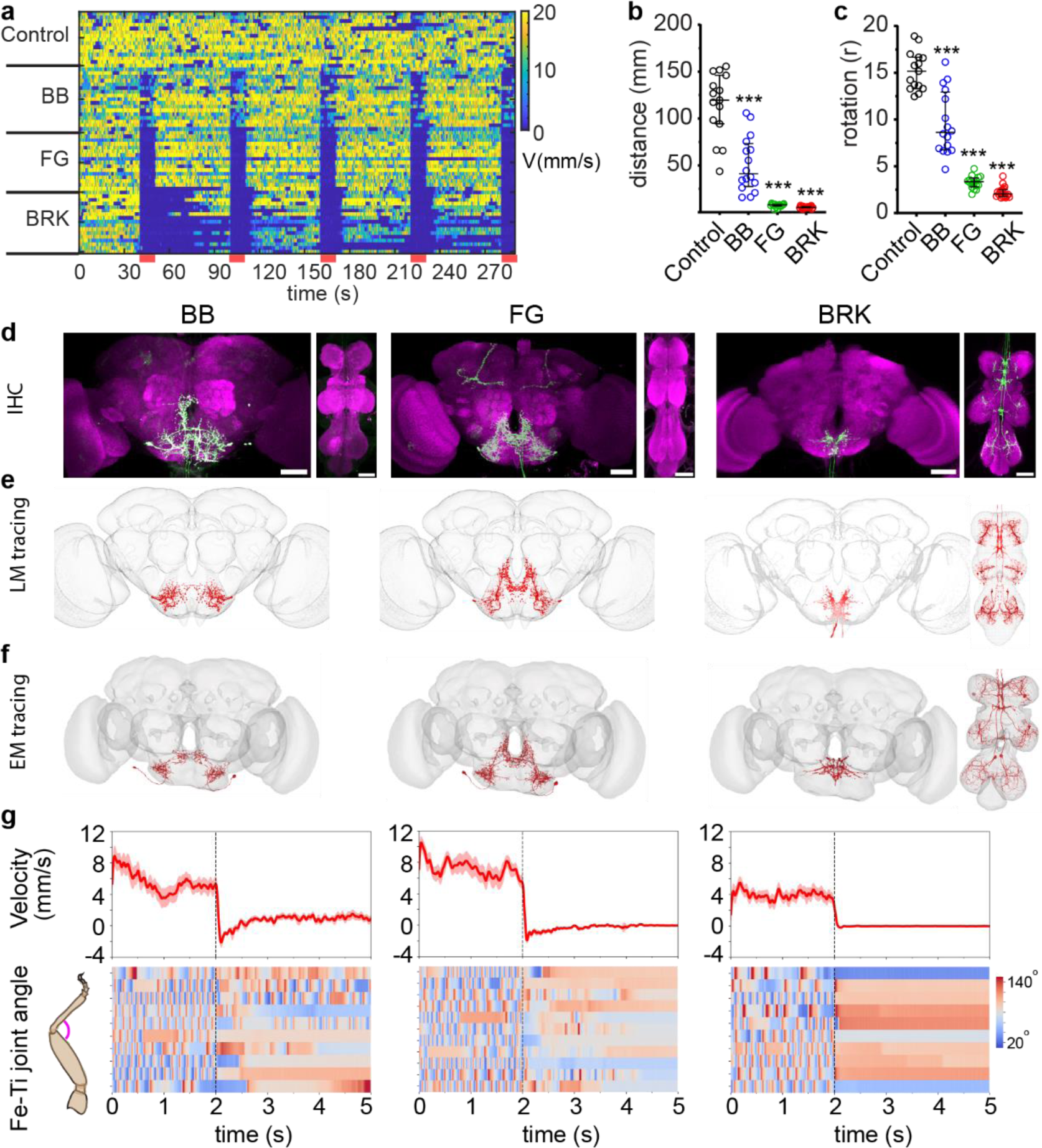
Identification of halting neurons: **a,b,c.** Translational velocity heatmap (**a**) (every row shows velocity of single fly, red bars indicate optogenetic stimulation), trial averaged travelled distance (**b**) and rotation (**c**) of free-walking control flies and flies expressing CsChrimson in halting neurons (BB, FG and BRK). n = 15-18 flies per genotype, non-parametric Mann-Whitney test compared to genetic-background control. (*** p<0.001). **d**,**e**,**f**. Immunohistochemistry (IHC) images of the selected genetic drivers (CsChrimson-mVenus:green, neuropil:magenta, scale-bar:50µm) (**d**), light microscopy based neuronal segmentation (**e**), electron microscopy based segmentation (**f**) of the BB (left), FG(middle) and BRK (right) halting neurons. **g.** Forward velocity (top) and femur-tibia joint angle heatmap (bottom, every row represents a different fly) of tethered walking flies during optogenetic stimulation of BB (left), FG (middle) and BRK(right) neurons. 3s stimulation starts at dotted vertical line in velocity plots. Refer to Supplementary Table 1 for full experimental genotypes and exact sample sizes.

Two of these neurons were previously morphologically described and were named as “foxglove” (FG) and “bluebell” (BB) for their characteristic arborizations resembling the respective flowers^33^ (Fig. 1d). Both FG and BB are SEZ local neurons that stochastically have a descending axonal projection which ends right after it enters the nerve cord. The FG line has weak ectopic expression in the Mushroom Body neuropil (Kenyon Cells) that does not contribute to halting phenotype (Extended Data Fig. 1b,c) The BB line also labels an additional cell-type (often unilaterally) anatomically described and named as “bluebird”^33^ (Fig. 1d). To disambiguate the contribution of FG, bluebell and bluebird, we also used another genetic reagent (SS31328) that specifically and reproducibly labels FG+BB without any off-targets, and drives robust halting on neural activation (Extended Data Fig. 1b,c). Finally, we discovered a previously unknown cell-type that drove the strongest halting phenotype (Fig. 1a-c, Extended Fig.1d). We named this the “brake” (BRK) neuron for the characteristic halting mechanism deployed by this neuron described in following sections. BRK neuronal type comprises six ascending neurons with their soma in each of the six leg-segment specific neuromeres of the ventral nerve cord (VNC) and axonal projections to the SEZ region of the brain (Figure 1d-f and Extended Data Fig. 2a,b). After identifying these neurons in the brain^5,6^ and nerve-cord^30–32^ connectome data, we found that although they have six distinct dendritic fields in the respective leg neuromeres, they have two shared output-zones, one in the tectulum area^34^ of the VNC and other in the SEZ region of the brain (Extended Data Fig. 2a,b). By stochastically activating subsets of BRK neurons, we found that activation of any of the BRK neurons is sufficient to drive halting (Extended Data Fig. 2c-e). We also generated segment specific BRK targeting drivers that further confirmed this observation (Extended Data Fig. 1e-f). This corroborated the connectome analysis which shows that all the six BRK neurons share common major outputs in both the brain and the VNC (Extended Data Fig. 2b).

This analysis of walking velocities (Fig. 1a-c) hinted at potential differences in the halting phenotypes induced by the three neuronal types (BB, FG and BRK). To precisely quantify such differences, we developed a pipeline for high-resolution three-dimensional leg kinematics analysis (Extended Data Fig. 3a,b) using pose-tracking (DeepLabCut^35^) followed by multi-camera-view based 3D joint-position reconstruction (Anipose^36^). Optogenetically activating the halting neurons in this assay revealed stark differences in the three halting phenotypes (Supplementary Video 1). BRK drove low-latency, long-duration halting bouts (Extended Data Fig. 3c) and joint angles locked in a specific position for the entire halt duration (Fig. 1g, Extended Data Fig. 3e). In contrast, FG and BB induced halts were interspersed with leg movements. The FG phenotype consisted of long halting bouts like BRK but still allowed brief repositioning of legs or grooming-like leg movements (Fig. 1g, Extended Data Fig. 3e). BB induced halts were much weaker comprising of stationary periods interspersed with slow walking periods and allowed more leg movements (Fig. 1g, Supplementary Movie 1). Unlike BRK, both FG and BB often allowed a few steps to be taken before the fly came to a halt (Extended Data Fig. 3d).

Identifying genetically targetable halting-promoting neurons driving distinct types of halting allowed us to ask if and how these distinct halting pathways interact with the walking circuits to promote halting.

### Specificity and hierarchy in Walk-Halt interactions

To address interactions between halting and walking pathways, we leveraged genetic tools targeting walking promoting neurons characterized in our previous work, viz. P9 that drives forward walking with turning^24,26^, BPN (Bolt Protocerebral Neurons) that drives straight forward walking^26^ and MDN (Moonwalker Descending Neurons) that drives backward walking^37^. We combined the split-Gal4 reagents in order to co-target pairs of “walk” and “halt” neurons in the same fly and tested the optogenetic co-activation phenotype. We confirmed that there was CsChrimson-mVenus expression in the desired neurons with little to no ectopic expression in the combined split-Gal4 experimental flies (Extended Data Fig. 4a). We co-activated flies expressing CsChrimson in all combinations of the three “walk” neurons (P9, BPN, MDN) and the three “halt” neurons (BRK, FG, BB, and the FG+BB line) and quantified their translational (Fig. 2a-c) and angular (Fig. 2d-f) velocities. Activation of BRK neurons overrode all walking commands (forward, backward and turning) when co-activated with the respective “walk” neurons, suggesting that they employ a dominant halting mechanism (Fig. 2, red traces show velocities falling to zero). On the other hand, FG and BB activation affected the “walk” neuron phenotypes to different degrees providing evidence for specificity and hierarchy in the way they interact with walking pathways.

**Fig. 2:**
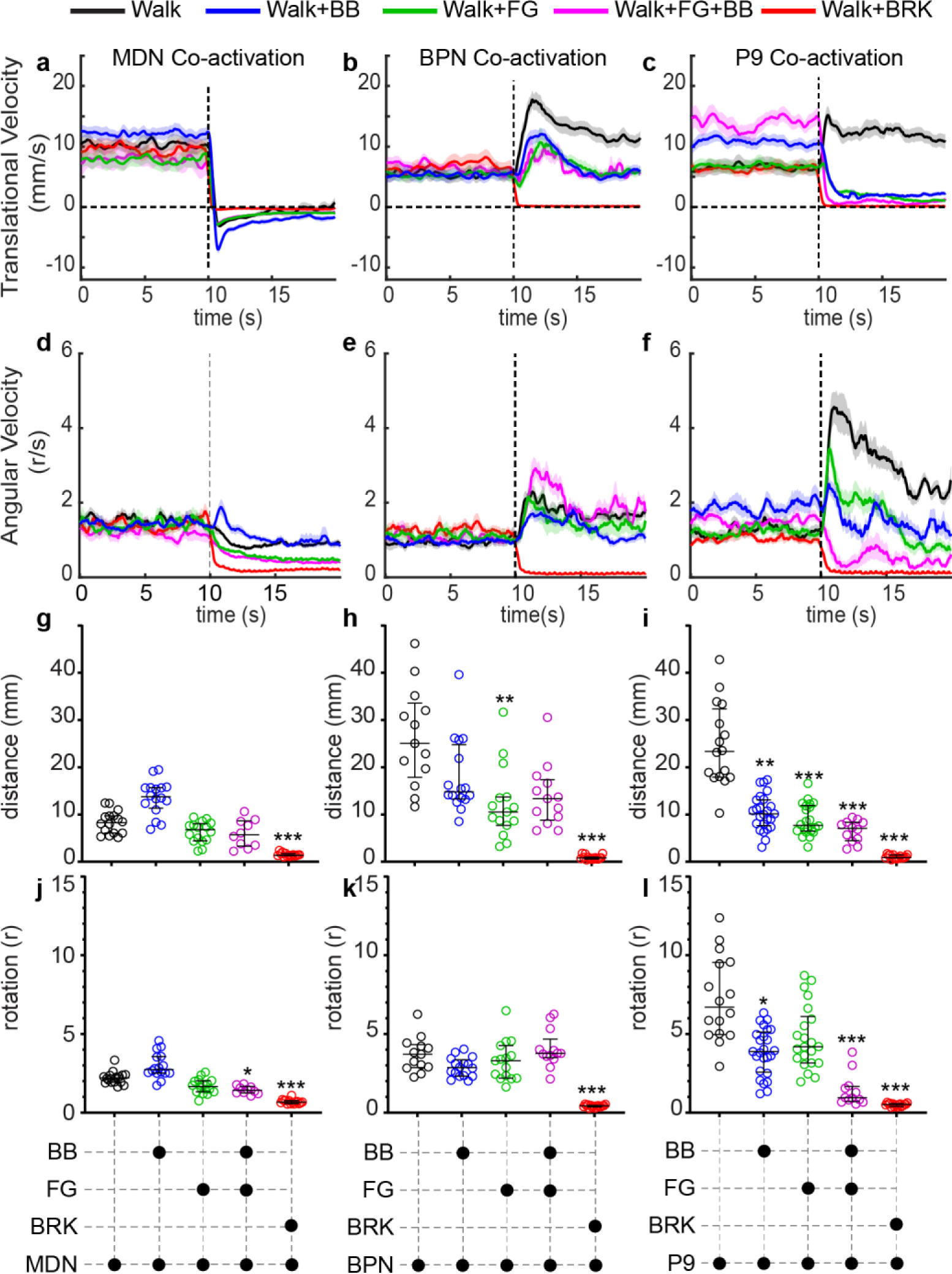
Walk-Halt Interactions: **a-i.** Trial averaged translational velocity (**a-c**), angular velocity (**d-f**), distance covered (**g-i**) and rotation (**j-l**) of free-walking flies with pairs of walk and halt neurons optogenetically stimulated. 10s stimulation starts at dotted vertical line in velocity plots. Each graph represents quantification for MDN (**a,d,g,j**), BPN(**b,e,h,k**) and P9 (**c,f,i,l**) co-activation experiments. n= 10-25 flies per genotype, Kruskal-Wallis test followed by Dunn’s multiple comparison with genetic-background control (*** p <0.001, **p<0.01, * p< 0.05). Supplementary Table 1 shows full experimental genotypes and exact sample sizes.

When the moonwalker neuron (MDN) is activated by itself, as reported in our previous work^37^, it does not drive a sustained backward walking bout. Instead, after about two seconds of continuous stimulation, the fly’s internal drive for forward walking often overcomes MDN driven backward walking, making the fly oscillate between backward and forward walking. This aspect of the phenotype is not clear in trial averaged plots (Fig. 2a) but apparent when we plot individual trial velocities (Extended Data Fig. 4b). MDN+FG co-activated flies completely lacked any forward walking (even in the later part of the trial), while backward walking remained unaffected. MDN+BB co-activated flies also showed a similar phenotype with the exception of a few forward walking events (Extended Data Fig. 4b). Just like MDN, other “walk” neurons also show a decreased phenotype when stimulated for longer than two seconds. So we restricted statistical comparison of distance (Fig. 2g-i) and rotation (Fig. 2j-l) covered by the fly to within two seconds of optogenetic stimulation. In case of MDN experiments, this analysis reflected distance and rotation of the fly in a backward walking state and showed that FG and BB have no effect on MDN driven backward walking. We conclude that unlike BRK, FG and BB are specifically forward walking suppression neurons.

When FG or BB was co-activated with BPN, the flies still showed slightly enhanced forward walking during optogenetic stimulation (Fig. 2b). While this increase in forward walking tended to be smaller than that of activation of BPN alone, only in case of FG could we see a statistically significant difference in the distance quantification (Fig. 2h). While the BPN+FG+BB co-activation showed decreased forward walking it was interspersed with intermittent jumps possibly due to ectopic expression in the VNC (Extended Data Fig. 4a) rendering the distance quantification unreliable (Fig. 2h). Based on these results, we conclude that FG can partially suppress the BPN driven forward walking.

On the contrary, when FG and/or BB were co-activated with P9, the flies showed a marked reduction in forward walking at stimulation onset (Fig. 2c, 2i). This is in stark contrast to increased forward walking observed when P9 is activated by itself (Fig. 2c). While BB+P9 or FG+BB+P9 co-activated flies showed a corresponding decrease in angular velocity compared to P9 stimulation (Fig. 2f, 2l), there was a major anomaly in case of FG+P9 co-activated flies. Despite showing a strong reduction in forward velocity, these flies still showed an increase in angular velocity on stimulation. While this effect is less evident in the averaged trial response plots (Fig. 2f), it was apparent when we quantified the rotation of the flies in the first two seconds after stimulation onset (Fig 2l). After observing the videos closely, we found that while the BB+P9 or BB+FG+P9 flies were predominantly halting during the optogenetic stimulation, FG+P9 flies were pivoting on the spot (pivoting quantified in Extended Data Fig. 4c, Supplementary Video 2). These observations revealed an unexpected modularity in the P9 downstream pathway: it shows that the P9 downstream pathway is segregated between forward walking and turning pathways. Since the FG+BB line has a similar phenotype as the BB line, we conclude that the ectopic neurons labeled in the BB line (bluebird) are not responsible for this phenotype. Taken together, we conclude that FG specifically overrides the forward walking component while BB overcomes both forward and turning components of the P9 driven neural pathways.

We conclude from these behavioral experiments, that the three halting pathways (BB, FG and BRK) indeed deploy different mechanisms to override the walking promotion pathways and impinge on walking circuit with specificity and hierarchy.

### The “Walk-OFF” mechanism: Manifesting halting by inhibition of critical walking promotion nodes in the SEZ region of the brain

Guided by the whole fly brain connectome and the FlyWire toolkit^38^, we set out to uncover the nodes where the halting and walking pathways converge. We first began by identifying, annotating and proofreading all the “walk” and “halt” neuronal types and their downstream partners in the connectome (Extended Data Fig. 5, Supplemental Table 3). This effort was complemented by the larger FlyWire consortium team^5^. We noticed that although P9 and MDN are cholinergic descending neurons (DNs: i.e. neurons that project from the brain to the nerve cord), they recruit other DNs before they exit the brain. While decapitated MDN-activated flies still show backward walking^39^, decapitated P9-activated flies do not (Extended Data Fig. 6j). This implies that the other DNs recruited by P9 (Extended Data Fig. 6f) are instrumental to its role in bringing about forward walking with turning. This is further corroborated by the fact that some of the P9 downstream DNs (viz. DNa01 and DNa02) have been previously implicated in ipsilateral forward turning^17,40^. BPN is a higher brain cholinergic neuronal type^26^ that directly recruits other DNs. Particularly, four previously uncharacterized DNs are strongly and directly connected to BPNs and we named these as BDN1, BDN2, BDN3 and BDN4 (Bolt Descending Neurons 1-4 respectively, Extended Data Fig. 6f). We therefore focused on identifying if any among these P9 and BPN pathway DNs are impinged upon by the halting pathways.

To do a comprehensive and unbiased analysis of this interaction between halting and walking pathways, we used a connectome-constrained whole-brain spiking-network model to reproduce the behavioral coactivation experiments *in-silico*. The model, described in detail here^4^, uses the entire connectivity matrix from the FlyWire data^5,38^, and assigns positive or negative weight to the connection based on the number of synapses and predicted neurotransmitter profile^41^. We further experimentally validated the neurotransmitter predictions for the three “halt” neurons using either FISH (Fluorescent In-situ Hybridization)^42^ or functional connectivity. Using these methods, we validate that BRK is an excitatory (potentially cholinergic) neuronal type, whereas FG and BB are GABAergic inhibitory neurons (Extended Data Fig. 6a-e).

In our model, the “walk” neuron was activated for the entire duration of the trial (1s) and the “halt” neuron was simultaneously activated for the central half of the trial (30 trials). An averaged trial response of the top 100 responding neurons is depicted in the firing rate heat-maps (Fig. 3a-d and Fig 3e-h). These heat-maps show that all three halting neurons inhibit different, partially overlapping subsets of neurons recruited by the “walk” pathways. Focusing on BB and FG results, these heat-maps indicate that while the P9 pathway is more broadly and strongly inhibited by BB, the BPN pathway is more broadly and strongly inhibited by FG. This is also apparent when we plot the inhibition (change in firing rate caused by the halt neuron) each neuron receives during the co-activation simulations with BB versus FG (Extended Data Fig. 6g-h), and aligns with the behavioral phenotypes observed in Fig 2.

**Fig. 3:**
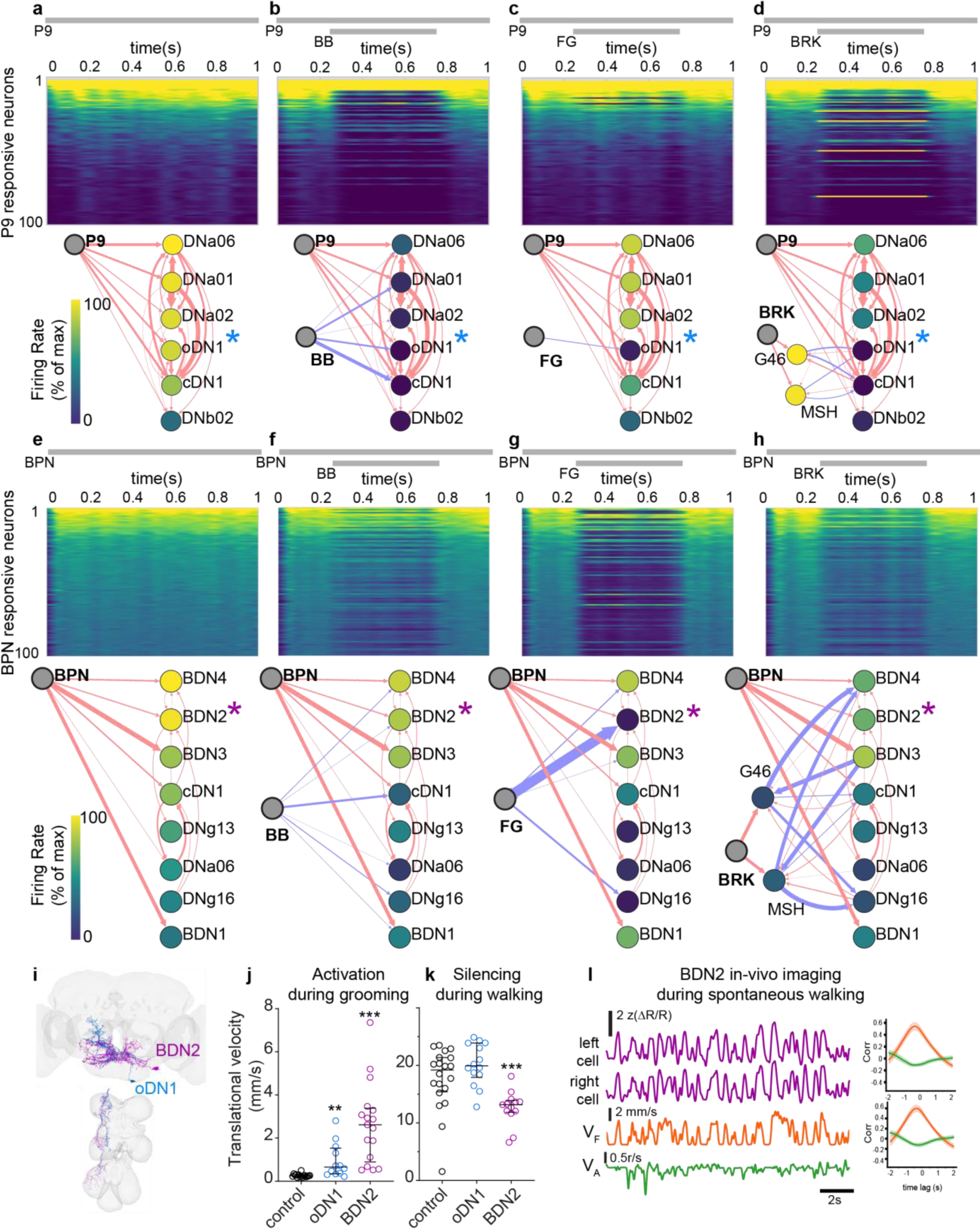
Walk-OFF mechanism: Modeling and functional data uncover critical walking promotion nodes inhibited by halting pathways: **a-h.** Simulation results for activating P9 (**a**), P9+BB (**b**), P9+FG (**c**), P9+BRK (**d**), BPN (**e**), BPN+BB (**f**), BPN+FG (**g**), BPN+BRK (**h**), activation paradigm indicated with grey bars on top. Trial averaged firing rate heatmap of top 100 P9 responding neurons (**a-d**) or top 100 BPN responding neurons (**e-h**) is shown on top, and connectome based wiring diagram for strongly recruited descending neurons is shown on bottom. Nodes in the wiring diagram are color coded based on firing rate in the full simulation, identical to the heatmap, color-scale is normalized to the max trial averaged firing rate achieved in the case of the respective walk neuron activation simulation. Red arrows indicate predicted excitatory connections and blue arrow indicates predicted inhibitory connections, arrow width corresponds to number of synapses and scales from 5 (thinnest) to 400 (thickest) synapses. Asterisks indicate neurons selected for further analysis in panels i-l. **i.** EM segmentation of oDN1 and BDN2. **j.** Translational velocity of stationary grooming flies induced to walk on optogenetic activation of oDN1 or BDN2 neurons. n=12-17 flies per genotype, non-parametric Mann-Whitney test compared to genetic-background control (** p<0.01, *** p<0.001). **k.** Translational velocity of free-walking flies with oDN1 or BDN2 optogenetic silenced using GtACR1. n= 13-20 flies per genotype, non-parametric Mann-Whitney test compared to genetic-background control (*** p<0.001). **l.** BDN2 activity (change in GCaMP6s/tdTomato reported as ΔR/R, top two traces corresponding to left and right BDN2 respectively) in a tethered fly spontaneously walking with forward velocity (V_F_) and angular velocity (V_A_), bottom traces. Right panels show cross-correlation of pooled data across 3 flies for activity of BDN2 left cell (top) and BDN2 right cell (bottom) with V_F_ (orange) and V_A_ (green). Supplementary Table 1 shows full experimental genotypes and exact sample sizes.

We next focused on the DNs strongly recruited by the walk pathways and generated synapse weighted graphs to simultaneously visualize the connectivity and firing rate (Fig. 3a-h bottom, firing rate coded neuroanatomy depicted in Supplementary Videos 3 and 4). In these plots (Fig. 3a-h bottom), the arrows (weighted by synapse numbers) represent direct connections between the nodes and the color of the node is determined by firing rate generated in the whole connectome network model. Therefore, the strongest direct connections (thick arrows) don’t always lead to strongest firing rates (yellow-colored nodes), indicating that indirect connections not shown in these graphs also impact the firing rate. In the case of P9, FG strongly and specifically inhibited a single DN named oDN1 that directly connected to both P9 and FG with high synapse numbers (Fig.3c, Supplementary Video 3). Comparing this with the behavioral phenotype (Fig. 2c,f,i,l) we infer that oDN1 is mainly relevant for driving translational velocity but not rotational velocity. On the contrary, the *in-silico* P9+BB co-activation showed that BB inhibits majority of DNs recruited by P9 (Fig. 3b, Supplementary Video 3) including oDN1 as well as previously implicated turning DNs (DNa01 and DNa02). This supports the results from behavioral experiments where we observed reduction of both translation and rotation in P9+BB or PB+FG+BB co-activation experiments (Fig. 2c,f,i,l).

In the case of DNs recruited by BPN, neither FG, nor BB inhibits the whole population (Fig 3e-g, Supplementary Video 4). This could explain why we never saw a complete loss of translation in the BPN coactivation behavioral experiments (Fig. 2). To probe why FG has a stronger effect in behavioral experiments, we compared the neurons that are differentially affected by FG versus BB. BDN2 stood apart as a DN specifically and strongly inhibited by FG (Figure 3g, Supplementary Video 4 and Extended Data Fig. 6g).

Taken together, the modeling and experimental data show that FG and BB are positioned to directly inhibit critical nodes in the P9 and BPN walking promotion pathways and decrease forward and/or rotational walking velocity. We term this the “walk-OFF” mechanism. Furthermore, oDN1 and BDN2 (Fig. 3i) stood apart as nodes that are strongly inhibited for reducing forward walking (Fig. 3a-h, Extended Data Fig. 6g,h). We therefore generated split-Gal4 drivers (Extended Data Fig. 6i) to specifically target these neurons and found that optogenetically activating oDN1 and BDN2 in non-walking flies (flies are covered with powder to make them stay in one place grooming themselves) drives walking initiation, with BDN2 showing a much stronger phenotype (Fig. 3j). Moreover, even decapitated oDN1 and BDN2 flies were able to initiate robust forward walking (Extended Data Fig. 6j) suggesting that unlike P9, the nerve cord outputs of these DNs recruit the downstream neural circuits sufficient for generating coordinated forward walking. When we optogenetically silenced these neurons using GtACR1^43^ in free-walking flies, only BDN2-silenced flies showed a reduction in forward velocity (Fig. 3k). This suggested that BDN2 might be critical during spontaneous walking, whereas oDN1 could be redundant or recruited only in specific contexts. To further confirm this, we imaged BDN2 activity (GCaMP) in spontaneously walking flies and found that its activity is strongly correlated to forward velocity of the fly, but not to angular velocity (Fig. 3l, Supplementary Video 5). Since BDN2 is directly and strongly inhibited by FG and not by BB, it could explain why FG drives stronger halting in spontaneously walking flies (Fig. 1).

*In-silico* coactivating the ascending outputs of BRK neurons along with P9 (Fig. 3d) and BPN (Fig. 3h) showed that while BRK is a cholinergic excitatory neuron, it recruits other predicted GABAergic inhibitory neurons in the SEZ which are poised to inhibit walking promotion DNs. In case of P9, BRK inhibits many of the critical walking-promotion DNs, including oDN1. In case of coactivation with BPN on the other hand, BRK inhibits a subset of DNs, with BDN2 only weakly inhibited in this simulation. This is in stark contrast to the complete absence of walking observed in the behavioral experiments when BRK is coactivated with any of the “walk” neurons (Fig. 2). Crucially, MDN, whose descending output in the nerve cord is sufficient to drive backward walking in decapitated flies, is also completely suppressed by BRK in behavioral experiments (Fig. 2a). As described earlier, BRK has two output regions, one in the nerve cord and one in the brain (Extended Data Fig. 2a,b). This suggests that, in addition to inhibiting some walking-promoting pathways in the brain akin to the “walk-OFF” mechanism, BRK could be directly acting in the nerve cord to bring about halting.

### The “Brake” mechanism: A ventral nerve cord specific mechanism to promote halting by increasing resistance at leg joints

To look further into the BRK driven halting, we first wondered if BRK neurons could cause the legs to “freeze” in any position like previously reported studies in mice^2^. We therefore re-visited the high-resolution 3D joint angle data from BRK optogenetic stimulation experiments. We found that unlike the recent findings in mice^2^, if the BRK optogenetic stimulation begins when the fly’s leg is in mid-air (during “swing phase” of walking), then the leg finishes the step in an unaltered manner, but freezes as soon as it lands on the ground (Fig. 4a). However, if the optogenetic stimulation begins while leg is in contact with the ground (“stance phase” of walking), then the leg stays locked in place at the onset of the stimulation (Fig. 4a). This is in contrast to FG stimulation where we see the leg always comes to stop, sometimes after taking a few steps, in a relaxed stance position, regardless of when the optogenetic stimulus started (Fig. 4b). This is clearly seen when we plot the probability density of the front leg femur-tibia joint angle distribution for the two stimulation-timing based conditions. The swing phase stimulation distribution shows that for majority of the BRK stimulation trials, the legs are locked in a position with the femur-tibia joint extended (Fig. 4c), which is how the front leg Fe-Ti joint looks at the end of a typical swing phase during forward walking. On the contrary, the stance phase BRK stimulation distribution shows a broader curve covering both acute as well as obtuse angles (Fig. 4c), showing that legs can get locked in different positions if they are already in the stance phase. A similar result is obtained when we analyze data for other leg-joints suggesting it is not a joint-specific phenomenon (Extended Data Fig. 7a, note that tibia-tarsus joint in any walking fly, like in Extended Data Fig. 3b, doesn’t change much during the stance phase and therefore shows smaller differences in the two test conditions). In line with this, BRK stimulation during swing phase does not alter the swing duration when compared to pre-stimulus swing phase duration (Fig. 4d).

**Fig. 4:**
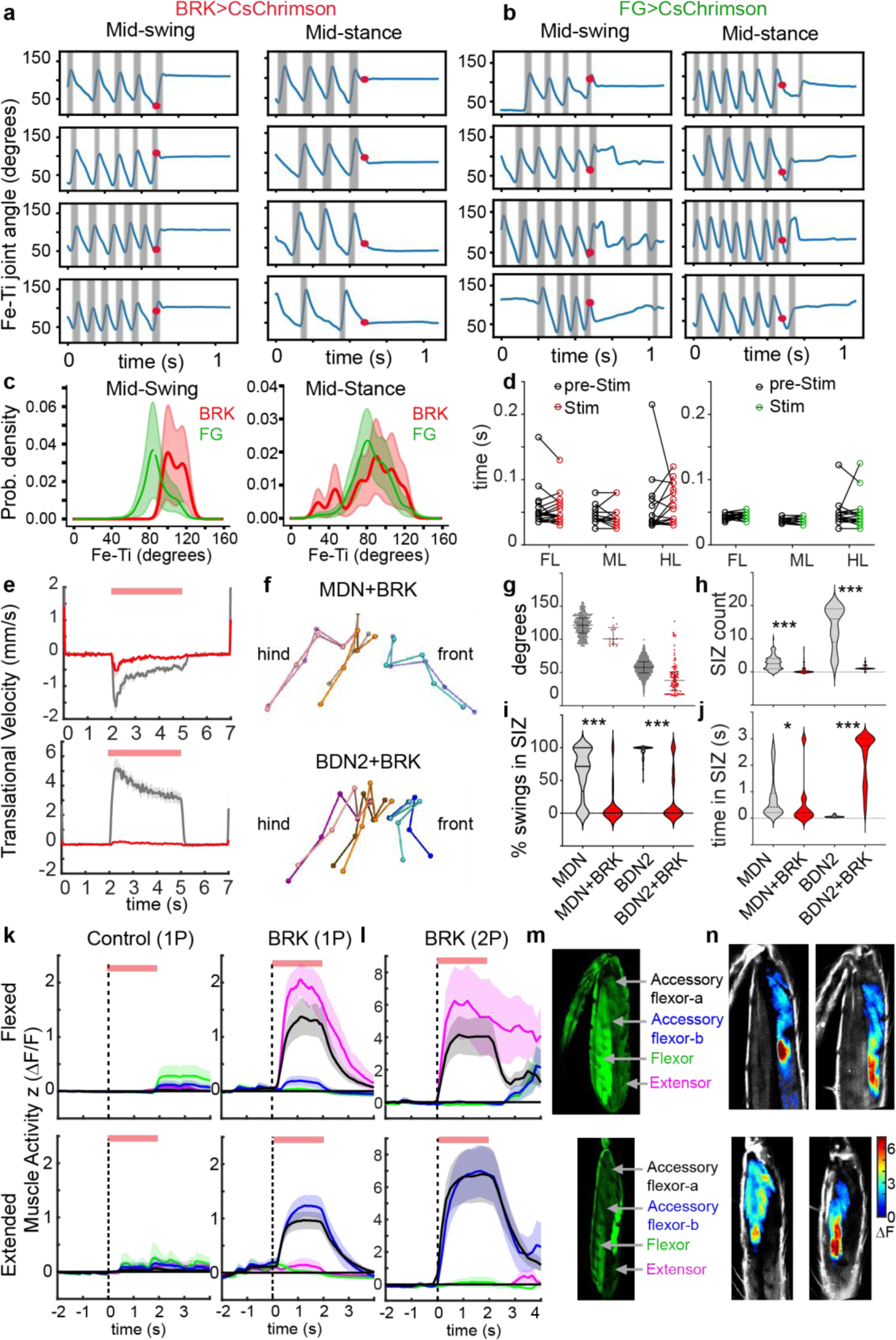
Break mechanism: VNC specific halting pathway for increasing resistance to leg movements. **a,b.** Data from 4 example flies (each row is one fly) showing front leg femur-tibia (Fe-Ti) joint angle position during optogenetic stimulation of BRK (**a**) or FG (**b**) in tethered walking flies. Left panels show cases where stimulation onset (indicated by red dot, stim lasts for 3s) was during the swing phase (grey shaded region) of walking, right panels show cases where stimulation onset (red dot) was during the stance phase of walking. **c**. Probability density of front leg Fe-Ti joint angle if BRK (red) or FG (green) was optogenetically activated during swing phase (left) or during stance phase (right)**. d.** Swing duration of BRK (left) or FG (right) swing-phase stimulated flies. Comparison between median swing-phase before stimulation and the swing-phase during which the stimulation occurred for front-legs (FL), mid-legs (ML) and hind legs (HL). n= 13-21 flies per genotype, Multiple paired t-test between pre-Stim and Stim showed no difference for all genotypes and legs. **e**. Averaged trial translational velocity of tethered decapitated flies while co-activating BRK with MDN (top) or BDN2 (bottom) n= 7-11 flies and 10 trials per fly, traces show mean±sem, top red bar indicates optogenetic stimulation. **f**. Video frame from 3D reconstructed tracking video showing front legs stuck in extended position (top) or flexed position (bottom) because of BRK+MDN or BRK+BDN2 stimulation respectively. **g**. Front leg Fe-Ti joint angle values at the onset of swing initiation in the respective genotypes, median±interquartile (MDN: 359 swings, MDN+BRK: 13 swings, BDN2: 2438 swings, BDN2+BRK: 115 swings). **h**. Number of times per trial when front leg Fe-Ti joint angle value entered Swing Initiation Zone (SIZ). **i.** Percent swings initiated after the front leg Fe-Ti joint entered SIZ. **j.** Time spent by the leg with Fe-Ti joint angle values in SIZ. h-j: Mann-Whitney comparison between walk and walk+BRK condition (exact n in Supplementary table 1, *** p<0.001, ** p <0.01) **k**. Left front leg femoral muscle activity (MHC>GCaMP6s, color coding as in m) while Fe-Ti joint is restrained in a flexed (top) or extended (bottom) position in control (left) or BRK>CsChrimson (right) flies during optogenetic stimulation (red bar) measured under epifluorescence (1P). mean±SEM, n= 6-7 flies per group (flexed or extended) and genotype **l.** Muscle activity measured with two-photon imaging (2P) in BRK>CsChrimson flies. mean±SEM, n= 6-7 flies per group (flexed or extended) and genotype. **m.** Anatomy of left front leg femoral muscles with Fe-Ti flexed (top) or extended (bottom) as seen in a confocal image of MHC>GCaMP6s flies depicting the nomenclature of muscles and color-coding of traces in k and l. **n.** Averaged trial ΔF image of 2 example BRK>CsChrimson flies with Fe-Ti joint flexed (top) or extended (bottom) showing characteristic resistance-reflex type activity profile observed on BRK stimulation during 2P imaging. Supplementary Table 1 shows full experimental genotypes and exact sample sizes.

This locking of joints during stance phase shows that the BRK phenotype is unlikely to be caused by an overall loss of muscle tone as it has been observed in some halting phenotypes^3^, but it rather suggests an active mechanism that engages specific muscles. But, how do the BRK neurons generate this stepping-cycle-phase dependent effect? We hypothesized that this could be driven by BRK nerve cord outputs upregulating the postural resistance reflex at the leg joints. The classical term “resistance reflex” is used to describe postural reflexes in all legged animals that allows them to maintain posture while they are in a stationary state^44,45^, e.g. if an animal is standing still and gets pushed from the back, then the resistance reflex kicks in and builds up muscle activity that resists the passively induced motion. When an animal switches from a stationary state to an active walking state, there is a “reflex-reversal” phenomenon that leads to a switch from “resistance-reflex” that resists leg movements, to “assistance-reflex” that assists stepping movements. Since these reflexes predominantly act on load-bearing legs that are in contact with the substrate, if the BRK neurons were to upregulate the postural resistance reflexes, the effect would primarily manifest as halting during the stance phase. To specifically test this hypothesis, we devised a behavioral experiment to test the specific effect of BRK nerve-cord outputs on stepping. It was previously shown that MDN activation in decapitated flies can drive backward walking^39^. We tested if co-activation of MDN and BRK in decapitated flies could override this backward stepping. The VNC outputs of BRK neurons were indeed sufficient for overriding the MDN induced backward walking, confirming that a ventral nerve cord specific circuit is recruited by BRK to arrest leg movements (Fig 4e).

MDN activation primarily drives hind-leg backward steps, while mid and front legs somewhat passively step in response to the induced backward translation^39^. This provided a perfect case for observing if BRK drives resistance to these passively induced backward steps in front legs. A typical MDN induced backward stepping in front legs shows a cyclic extension, levation and flexion of the leg. Focusing on the femur-tibia (Fe-Ti) joint of the front leg, we observe that this joint reaches highly extended values before a swing is initiated (Fig. 4f,g). For quantification purpose, we refer to all Fe-Ti joint angle values above 109 degrees (75^th^ percentile of swing initiation Fe-Ti values) as the “swing –initiation-zone” (SIZ). When BRK is co-activated with MDN, the Fe-Ti angle remains locked in place and never reaches the SIZ in a large number of trials (Fig. 4h). However, in some cases the MDN activation causes the Fe-Ti joint to reach the SIZ (Fig. 4f) but, as is typical for resistance reflex, this rarely led to levation of the leg. The leg, despite being in an extended Fe-Ti position, still remained firmly perched on the substrate as indicated by decreased swing events in SIZ (Fig. 4f,i, Supplementary Video 6).

To perform a complementary analysis for forward walking, we repeated this set of experiments with BDN2 since it was identified as a critical component of forward walking and drove forward walking in decapitated flies. In contrast to front-leg stepping during backward walking (extension->levation->flexion), during forward walking (flexion->levation->extension) the SIZ for Fe-Ti joint lies in the flexed angle values (Fig. 4g, SIZ is Fe-Ti < 66 degrees, which corresponds to 25^th^ percentile of swing initiation Fe-Ti values). Despite this difference, co-activation with BRK interrupted forward walking (Fig. 4e, Extended Data Fig. 7b,c) and showed similar results with respect to leg kinematics. The legs rarely entered the SIZ (Fig. 4h), and if they entered the SIZ, they stayed firmly perched on the ball with Fe-Ti joint angle locked in highly flexed position (Fig 4f,i,j, Supplementary Video 6). Taken together, this showed that BRK activation led the flies to resist both extension and flexion at the Fe-Ti joint, and in cases where leg was (forcibly) flexed/extended, it still did not lead to initiation of swing phase.

To functionally confirm if BRK was driving a resistance state in the legs, we analyzed activity (GCaMP fluorescence, 1-photon wide-field imaging) of the front-leg femoral muscles while inducing a resistance state in the leg at the femur-tibia joint and simultaneously activating BRK. If we forcibly kept the femur-tibia joint in a flexed position and activated BRK, the tibia-extensors along with accessory flexors-a showed increased activity; differently, if we kept the leg in a fully extended position, BRK activation stimulated two groups of tibia accessory flexors (Fig. 4k, Supplementary Video 7). To get a clearer picture of these muscles, we repeated the BRK stimulation trials under a two-photon microscope and found that just like in 1P imaging, the accessory flexor-a is activated in both flexed and extended conditions. A second bundle of muscle fibers, which we named as accessory-flexor-b, was recruited only in the extended condition, while extensors were recruited in flexed condition (Fig 4l-n, Supplementary Video 7). By comparing the anatomy of these muscles (Fig. 4m-n) to published work using the same muscle driver^32,46,47^, we conclude that accessory-flexor-a corresponds to the muscles in the distal part of the femur innervated by slow motor neurons, also referred to as “reductors”. Our observation that BRK recruits these “slow” flexors in both flexed and extended condition fits with previous reports that have implicated these muscles in posture control and resistance reflex^47^. The accessory-flexor-b are more proximal and were not categorized as a different muscle in the previous studies. Given their location, it is likely that they are innervated by intermediate or fast motor neurons and could explain their specific recruitment in the extended position. We never observed the large fast flexors being recruited by BRK neurons, even though they were clear during spontaneous activity uncorrelated to optogenetic stimulus. We happened to record a few instances where there was large spontaneous rhythmic activity in fast flexors and extensors during the optogenetic stimulation. In such cases we saw that upon BRK stimulation, the pre-stimulus activity gets inhibited and is simultaneously replaced by the characteristic activity profile of the BRK stimulation based on the joint position (Supplementary Video 8). This implies that BRK is bringing about a concerted change that shapes the overall muscle activity profile towards a resistance reflex-like activity. We also noticed similar “slow-muscle/reductor-muscle” activity in other leg segments (data not shown) that could underlie similar joint angle phenotype observed at other joints (Extended Data Fig. 7a). Taken together, this shows that in restrained legs, BRK neurons recruit the postural control and resistance reflex implicated “slow” flexors and depending on the position of the joint also recruit either extensors (flexed position) or accessory flexor-b (extended position).

To get another view at the resistance reflex, we attached a metal pin to the tibia and actively moved it with a motorized magnet while imaging the muscles in the femur (based on setup in^48^) and providing continuous optogenetic stimulation. Here, both control and test flies showed a resistance reflex-like muscle activity and confirmed our expected recruitment of accessory-flexors and extensors (fast-flexors were not recruited even in this condition). However, the BRK stimulated flies showed both larger as well as more robust (less variance) responses compared to controls (Extended Data Fig. 7d).

Combining the results from behavioral and functional studies, we conclude that BRK drives a stance-phase specific resistance state at the leg joints via a nerve cord specific neural circuit.

### Task-specific recruitment of halting pathways

#### FG and BB pathways are recruited for halting in the context of feeding

We next addressed if and when does the animal actually use the FG, BB and BRK halting pathways. Mining the connectome for inputs to the halting neurons showed that FG and BB are downstream to previously characterized command-like feeding neurons (Fdg^49^). Fdg neurons respond to sucrose^50^ and activation of Fdg neurons drives halting followed by proboscis extension^49^. Indeed, activating the sweet sensory neurons drove increased activity in FG and BB both *in-vivo* as well as *in-silico* (Fig. 5a,b). Moreover, the degree to which the respective halting neurons were activated was also similar across experiments and the model, with FG showing a large response and BB showing smaller but sustained responses (Fig. 5b). We also noticed that FG responds to sugar sensory neurons in a state dependent manner with much stronger responses in starved flies compared to fed flies (Extended Data Fig. 8a). This could be because the sweet sensory pathway is upregulated by hunger signals at multiple points, as indicated by previous work^50^. To understand how different activity levels in the sweet sensory pathway could interact with different activity levels in the walking pathway, we simulated the network model, while titrating the levels of sweet sensory pathway and P9/BPN walk pathways simultaneously. We found that the important nodes, oDN1 and BDN2 in the respective walking promotion pathways (P9 and BPN) are tightly regulated as a function of activity levels in the sweet sensory pathway, showing that there is an excitation/inhibition balance at these nodes (Fig. 5c,d). To test if FG is critical for relaying the inhibition from the sweet sensory pathway to the walking nodes, we repeated the above simulation but while silencing FG in the model. This clearly shifted the excitation/inhibition balance at the walking nodes, such that we could now see release of inhibition (“disinhibition”) of oDN1 and BDN2 when compared to the results in the FG intact model (Fig. 5c,d). This effect was much stronger in case of BDN2 as compared to oDN1, such that when FG is silenced the activity profile of BDN2 is almost unaffected by sugar GRN activation. This implies that while FG is critical for inhibiting BDN2 in the context of feeding, there could be other pathways besides FG that drive the oDN1 inhibition. Since BDN2 is recruited during spontaneous walking and silencing BDN2 leads to decreased forward velocity, inhibition of BDN2 via FG could be relevant for halting in the context of foraging/feeding.

**Fig. 5:**
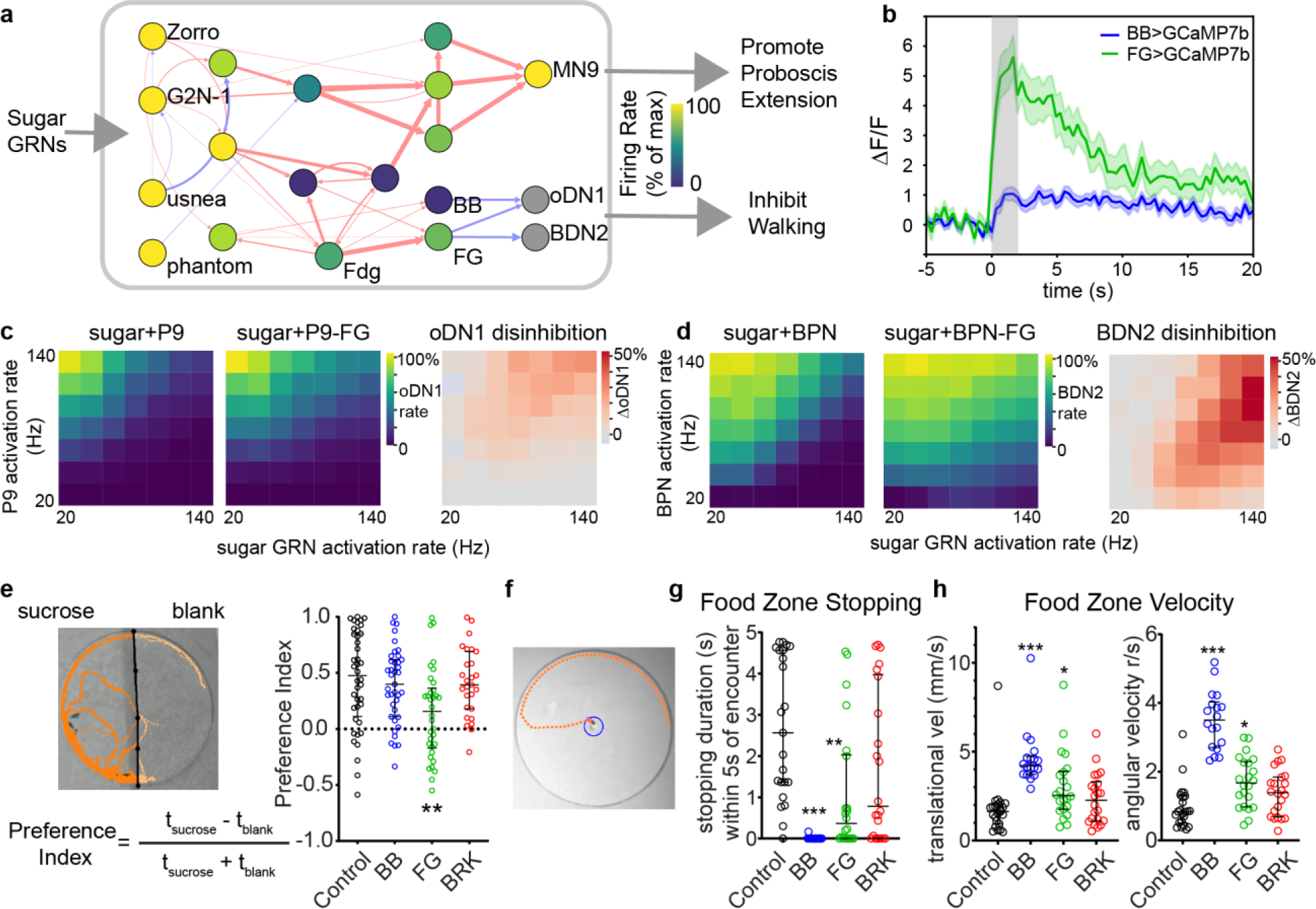
FG and BB recruited for halting in the context of feeding. **a.** Simulation of sucrose sensing pathway (sugar GRNs activated at 150Hz) with addition of FG and BB activity profile and showing inhibitory connections to oDN1 and BDN2. Nodes are color coded by firing rate and arrows weighted by synapse number as in Fig.3 **b**. FG or BB activity (*in-vivo* GCaMP7b imaging, mean±SEM) during optogenetic stimulation (grey background) of sweet sensory neurons (Gr5a), in starved flies. **c**. Simulation of oDN1 firing rate while co-stimulating P9 and sugar GRNs across range of stimulation rates in intact model (left) or with FG silenced model (middle). Difference between middle and left panel is depicted as the release of oDN1 inhibition in right panel. **d**. BDN2 firing rate while co-stimulating BPN and sugar GRNs across range of stimulation rates in intact model (left) or with FG silenced model (middle). Difference between middle and left panel is depicted as the release of oDN1 inhibition in right panel. c,d firing rates are normalized to max firing rate in left panel of c or d. **e.** Video frame with overlaid example fly-trajectory from two-choice assay (left) and preference index (right) of sucrose over blank, for flies with BB, FG, or BRK optogenetically silenced using GtACR1. n= 27-40 flies per genotype, Kruskal-Wallis test followed by Dunn’s comparison (** p<0.01). **f** Video frame with overlaid example fly-trajectory from food-blob interaction assay showing food interaction zone as a blue circle. **g**. Stopping bout duration within 5s of food encounter for flies with optogenetically silenced halting neurons (GtACR1). **h.** Translational (left) or angular (right) velocity of flies in the food zone within 5s of food encounter for flies with FG, BB or BRK silenced using GtACR1. g,h: n= 18-22 flies per genotype, Kruskal-Wallis test followed by Dunn’s comparison (*** p<0.001, ** p<0.01, *p<0.05). Supplementary Table 1 shows full experimental genotypes and exact sample sizes.

We next optogenetically silenced the halting neurons while providing the flies a choice between blank versus sucrose substrate (dried filter paper where flies can sense the sucrose but cannot consume it, assay inspired from^51^). While control flies or flies with BRK/BB silenced spent most of their time on the sucrose side, FG silenced showed a stark deficit in their ability to stay on the sucrose side (Fig. 5e). To test specifically whether flies use FG and/or BB for halting in the context of feeding, we analyzed their interaction with a small blob (3 mm diameter) of sucrose in agarose (200mM) placed at the center of a large circular arena (50 mm diameter, Fig. 5f). Both FG and BB silenced flies showed shorter stopping bouts near the sucrose blob compared to controls or BRK silenced flies, which showed prolonged feeding related stops (Fig. 5g). All flies showed some degree of slowing down when they first encountered the food blob. However, FG and BB silenced flies show increased velocities in close proximity of the blob compared to controls (Fig. 5h). Here, BB silenced flies showed a much stronger phenotype with a large increase in both translational and angular velocities. BB silenced flies had normal velocity profile pre-encounter of the food, but show a sudden increase in walking and especially turnin, post encounter (Extended Data Fig. 8c). This could imply a potential role for BB in constraining forward and angular velocities during foraging type behaviors that involve a local search following food encounter^52–54^.

These imaging, modeling and behavioral experiments, taken together, confirm that FG and BB are recruited for halting and locomotor control in the context of feeding. Notably, connectomics revealed that this halting pathway diverges from the previously described proboscis extension pathway necessary for extending the proboscis for food consumption^4,50^ (Fig. 5a). In agreement with this, FG/BB silencing had no effect on the proboscis extension response when presented with sucrose (Extended Data Fig. 8b).

#### BRK pathway is recruited for halting in the context of grooming

We did not find any previously characterized neurons directly upstream to BRK neurons in the VNC connectome. To understand when the animal recruits BRK neurons, we monitored the activity (GCaMP fluorescence) of these neurons while simultaneously recording both the posture, as well as walking velocities of tethered flies spontaneously walking/halting on an air-supported ball (Fig. 6a). We developed a preparation inspired by previous work^17^, where we could image a region of the ventral nerve cord that permits unambiguous identification of the axons belonging to each of the six BRK neurons (Fig. 6b). We first noticed that BRK neuronal activity only rises during a subset of halting events. By annotating the synchronously recorded behavior, we found that BRK activity correlated with either “ball-pushing” or “grooming” events (Fig. 6C). Since “ball pushing” is likely a special behavior that only happens in a tethered fly walking on an air-supported ball, we focused on the grooming events. BRK activity correlated in a segment-specific manner to the respective segment-specific grooming events. Most prominent among these were front leg rubs and hind leg rubs that coincided with large correlated activity bumps in foreleg and hind-leg specific BRK neurons as seen in pixel-wise correlation maps (Fig. 6d and Supplementary Video 9). Aligning BRK activity profile across 6 flies (56 front and 63 hind leg grooming events), it was clear that this grooming correlated rise in activity synchronized with a coincident dip in walking velocity (Fig 6g,h), suggesting that BRK neurons are recruited in order to initiate and maintain a stable halted state during grooming events. There were even a few rare mid-leg grooming events (one mid-leg+two hind-legs) that showed an expected correlation with mid-leg neuropil innervating BRK neurons (Fig. 6e,f). Since all BRK neurons share common outputs, based on BRK mechanism insights (Fig. 4) we suggest the following hypothesis: whenever a particular subset of BRK neurons is recruited during grooming, it stabilizes the legs that are already in stance position to promote and maintain a halted posture, without interfering with ongoing grooming or other unloaded leg movements.

**Fig. 6:**
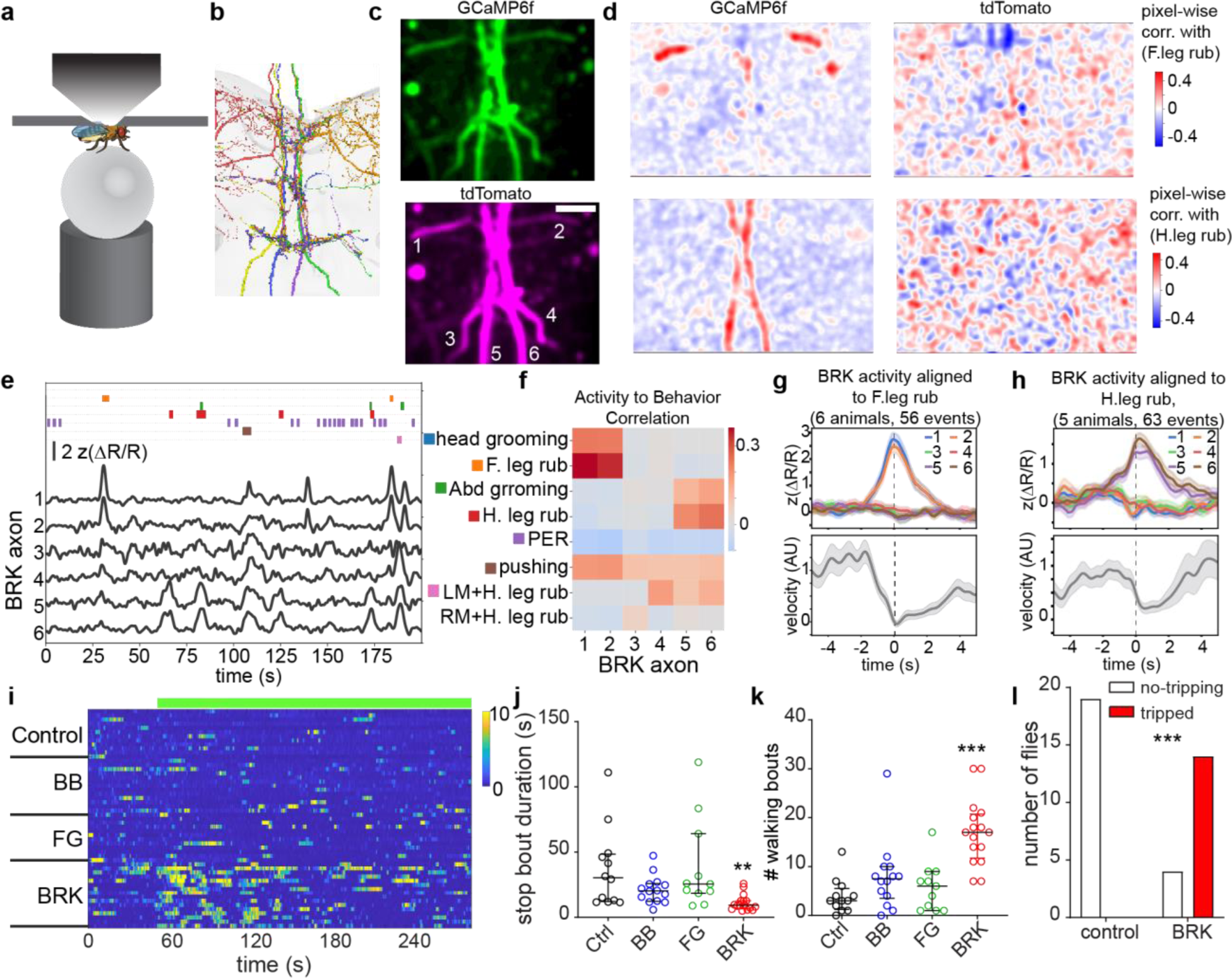
BRK is recruited for halting in the context of grooming. **a.** Schematic of *in-vivo* VNC imaging preparation. **b.** EM segmentation of BRK near the output zone in VNC tectulum region which was imaged in following panels. **c.** Example fly, averaged image of green (GCaMP6f) and magenta (tdTomato) channels with BRK neuron numbers 1-6 juxtaposed to the 6 axons from BRK neurons used as ROIs for quantification (1,2: front leg BRK, 3,4: mid-leg BRK, 5,6: hind leg BRK). **d.** Pixel-wise correlation map of the same example fly from c showing pixels with intensity correlated to front leg-rubbing events (top) or hind-leg rubbing events (bottom). GCaMP channel shows strong segment specific correlation to corresponding grooming events (left) which are absent in the tdTomato channel (right). **e.** Example imaging session that shows all ethogram (top) and z-scored ΔR/R (bottom) for each BRK neuron (neuron numbers as indicated in c.) **f.** Correlation between BRK activity and annotated behaviors (neuron numbers as in c) for pooled data across 6 flies. **g,h.** z-scored ΔR/R (top) aligned with respect to initiation of front leg rubs (**g**) or hind leg rubs (**h**), and corresponding forward velocity of the fly (bottom). **i-k.** Translational velocity heatmap (i), stop bout duration (j) and number of walking bouts (k) of powdered grooming induced flies during optogenetic silencing of control, BB, FG or BRK with GtACR1 (top green bar in i depicts optogenetic silencing) n= 11-16 flies per genotype, Kruskal-Wallis test followed by Dunn’s comparison (*** p<0.001, ** p<0.01). **l**. Number of decapitated, powdered flies that tripped versus remained stable while grooming, during optogenetic silencing of control or BRK with GtACR1, n= 18-19 flies per genotype, Fisher’s exact test (*** p< 0.001). Supplementary Table 1 shows full experimental genotypes and exact sample sizes.

To directly test this hypothesis, we optogenetically silenced BRK neurons in flies that were induced to groom by covering them with powder^55,56^. BRK optogenetic silencing did not completely abolish grooming, but led to interruption of ongoing grooming by intermittent walking events (Fig. 6i-k). On the contrary, control flies or flies with FG or BB silenced showed prolonged uninterrupted grooming (Fig. 6i-k). Upon close inspection of the video (Supplementary Video 10), it was apparent that on BRK silencing, the overall posture of the flies becomes destabilized during grooming. To probe more into this aspect and also address if this phenotype was due to the VNC outputs of BRK neurons, we repeated this experiment in decapitated flies. Consistent with literature^56^, control decapitated flies showed robust grooming when covered with powder. Since decapitated flies do not have an inherent forward walking drive like intact flies, it was unsurprising that we did not see any walking on BRK silencing. However, we saw that the powdered and decapitated BRK silenced flies often lost their balance and tripped-over showing that they lacked the postural control present in control flies (Fig. 6l). Close observation of the tripping events showed that this often happened as a result of extra legs lifted off the ground at the same time (Supplementary Video 9). Taken together, these *in-vivo* imaging and neuronal silencing experiments imply that BRK is recruited for proper halting and postural control in the context of grooming. To see if BRK could receive segment specific grooming relevant information, we mined the nerve cord connectome data^31,32^. Indeed, the six BRK neurons are well-positioned to receive segment specific leg mechanosensory and proporiosensory inputs relevant for grooming (Extended Data Fig. 9a).

We thus elucidate distinct behavioral contexts in which each of the halting pathways is recruited.

### Connectome guided prediction of novel halting pathways

Having identified the critical nodes in the walking pathways that are inhibited by the halting neurons, we wondered if we could use this information to gain insight into other halting pathways that could be potentially recruited in other behavioral contexts. Previously, we implicated an ascending neuron named MAN (Moonwalker Ascending Neuron) in the maintenance of backward walking by inhibition of forward walking. MAN activation using CsChrimson indeed drove halting (Extended Data Fig. 10a-b). We next modeled MAN coactivation with the other “walk” neurons and found that important forward walking nodes, including BDN2 and oDN1, get inhibited (Extended Data Fig. 10c-d). By mining the connectome, we uncovered a pathway leading from the predicted cholinergic MAN to BDN2 via previously uncharacterized GABAergic neurons (G129, CB283). We next looked at the well characterized egg-laying pathway, since egg-laying involves sustained halting in order to lay the eggs on the appropriate substrate. Moreover, the oviposition descending neurons (oviDNs) implicated as the “command-like” egg laying neurons^57^ have been recently shown to induce halting on optogenetic activation^58^. Indeed, when we activated oviDNs *in-silico*, it inhibited both oDN1 and BDN2 (Extended Data Fig 10c-d). Connectomics revealed the major inhibitory neuron recruited by oviDN is CB283. We therefore predict CB283 to be a halting neuron recruited in the context of egg-laying. Finally, mining for predicted inhibitory inputs to oDN1 and BDN2 revealed a list of potential halting neurons, many of which are also poised to inhibit other potential walking promotion DNs revealed in our previous simulations (Extended Data Fig. 10e). These predicted halting neurons receive input from different upstream neurons, and could underlie execution of halting in other behavioral contexts (fear, sleep, courtship etc.). However, major input zone of these predicted neurons is still in the SEZ region of the brain (GNG, VES neuropils which are part of the SEZ), implying their recruitment closer to the motor output, similar to FG and BB (Extended Data Fig. 10f,g).

## Discussion

Our work elucidates two fundamental halting mechanisms in *Drosophila*. The first mechanism, which we categorized as the “walk-OFF” mechanism, involves GABAergic neurons that inhibit walking promotion pathways. This could be compared to taking one’s foot off the gas-pedal of a car. Inhibition of action-promotion neurons is a cessation mechanism proposed in different behaviors across organisms^59–64^. Using a combination of genetic tools and connectomics, we could go beyond identification of this mechanism and gain major insights into the organization of the walking-control circuits in flies: a) We found that previously characterized walking promotion DNs (P9 and MDN) or local brain neurons (BPNs) recruit populations of heterogeneous DNs via their outputs in the SEZ area of the brain, suggesting that DN populations bring about a wide-spread modulation of nerve cord circuits for driving specific walking behaviors. b) Through identifying convergence points of halting and walking pathways, we uncovered the essential nodes in these DN populations (e.g. oDN1, BDN2) for walking initiation and maintenance. oDN1 and BDN2 are the first identified DNs in *Drosophila* that can initiate and maintain a coordinated forward-walking state even in decapitated animals suggesting that their outputs in the nerve cord recruit the essential pattern generation machinery. Following these pathways in the nerve cord will lead to identification of core elements of walking pattern generation that have been so far elusive in *Drosophila*. c) We show that different halting pathways inhibit different subsets of walking promotion neurons. Thus, unlike a car, a fruit fly’s brain has several “gas-pedals” (aka. walking promotion neurons) and different halting pathways are poised to inhibit different subsets of these “gas-pedals”. This led us to uncover that populations of turning and forward walking promoting descending neurons could be differentially inhibited by BB and FG halting pathways respectively (Supplementary Video 3). This shows a modularity in the descending control of forward walking and turning that is consistent with recent whole-brain imaging work that shows orthogonal neural activity profiles during turning versus forward walking in *Drosophila*^8^. Remarkably, this also aligns with modularity at the level of brainstem spinal projection neurons driving forward locomotion and turning observed in vertebrates (zebrafish^65^, mice ^20,66^). Vertebrates could therefore also have halting pathways that differentially inhibit forward locomotion and turning. Differential inhibition of forward walking versus turning could be instrumental in biasing the animal’s locomotor output towards local search (exploitation) versus long straight walking bouts (exploration), respectively and could underlie the fundamental “exploitation versus exploration” decisions all animals undertake during foraging^67–69^.

The second mechanism, which we term as the “brake mechanism”, brings about active resistance at the leg joints, much like the brakes of a car resisting motion of the wheels. Also, similar to how one releases the gas pedal while pressing on the brakes to avoid skidding, the newly identified “brake” neurons (BRK) inhibit the walking-promotion neurons while simultaneously promoting resistance in the legs, so that legs don’t end up in strained configurations. In line with this, when we removed the ability of BRK to inhibit walking-promotion neurons by decapitating the fly and co-activated BRK with walking-promoting neurons, we indeed pushed the fly into strained configurations (Fig. 4f, Supplementary Video 6). Furthermore, we used behavioral and muscle imaging experiments to show that BRK neurons only promote resistance in legs that are in stance position, thereby avoiding halting in unstable positions with legs in mid-air. BRK neurons achieve this by primarily recruiting the slow-type muscles that have been previously implicated in postural control and resistance reflex^47^. This also agrees with previous neuromechanical modeling work that implicated these muscles in promotion of stopping in stick-insects^70^. The pathway from BRK to muscle activation is likely gated by proprioceptive inputs (similar to previously described stick-insect proprioception-gated neurons^71^) that confers the stance-phase specific behavioral response (Extended Data Fig. 9b shows putative pathway from BRK to Acc. Flexors with proprioceptive feedback). Further work will be needed to uncover the exact nature of these transformations that lead to stepping-phase specific effect of BRK neurons. Moreover, we show that this mechanism of simultaneously inhibiting walking promotion neurons while increasing resistance at leg joints is a dominant halting mechanism that overrides all walking promotion signals. We speculate that this mechanism could underlie other halting pathways in flies, as well as analogous pathways in other animals (including mammals) for execution of mechanically stable halting in situations where it has to be dominant over conflicting walking commands. Our results encourage the search for analogous spinal ascending pathways in vertebrates that could be instrumental for dominant halting phenotypes.

We also found that different halting pathways are recruited in distinct behavioral contexts. While FG and BB pathways are recruited for halting during feeding and foraging, the BRK pathway is recruited during grooming. We cannot exclude that other contexts also recruit these pathways and we also predicted novel halting neurons that could be recruited in different contexts. Comparing across the connectome-informed circuits of all the experimentally verified or predicted halting pathways, we identified a common theme that suggests a generic neural circuit motif for halting. For promoting halting to allow other stationary behaviors like feeding, egg-laying or grooming, the halting pathway branches off from the behavior-specific sensory-motor pathway and impinges on core walking promoting circuits. The halting neurons, themselves need not encode for other core attributes of the behavioral motor pattern (e.g. proboscis extension in case of feeding, ovipositor extrusion in case of egg-laying, or grooming leg movements) but rather drive the initiation and maintenance of a halted state during the behavior. This halting branch is typically located downstream to higher order (“command-like”) neurons in the behavioral sensory-motor circuit, and closer to the motor output of the pathway. This emergence of modularity downstream in the circuit could be an efficient circuit organization for targeting similar population of walking promotion pathways in different contexts. It could also provide flexible recruitment of motor outputs while engaged in the same behavior (e.g. a fly need not extend its proboscis every time it halts during foraging, so the PER circuit and halting pathway could be decoupled when necessary, downstream of the command like “Fdg” neurons).

Despite this conserved motif shared across all halting pathways, the specific targets and mechanisms deployed by the halting neurons to promote halting are distinct. We speculate that it is this distinction that confers the hierarchy underlying how halting versus walking decisions are executed in a context appropriate manner.

## Methods

### Experimental animals

We used *Drosophila melanogaster* raised on standard cornmeal-agar medium supplemented with baker’s yeast and incubated at 25°C with 60% humidity and 12h light/dark cycle throughout development and adulthood unless otherwise stated. The optogenetic manipulation experimental flies were collected on retinal food and again transferred to fresh retinal food 1-2 days prior to testing. Retinal food contains standard fly food with freshly added 400μM all-trans retinal (Sigma-Aldrich R2500). These flies were kept in the dark for entire life-cycle until tested. The age and sex of animals tested is indicated in the method sections below. All full experimental genotypes, exact sample size per genotype and source of the genetic reagents are described in Supplementary Table 1 and Supplementary Table 2.

### Identification and generation of halt neuron specific drivers

The neural activation screen comprised of an extended version of our previously published work^26^ with added lines from the SEZ-splitGal4 collection^33^. While in our previous work^26^ we focused on lines that increased walking on optogenetic stimulation, here we focus on lines that decreased walking. We could narrow the set of interesting lines to 11 drivers based on their locomotor phenotype and expression levels. Among the SEZ lines, we focused on 3 lines where we could unambiguously identify the neurons both in the light microscopy^27^ as well as electron microscopy^5,31,32,72^ datasets. Additionally, we found three Gal4 drivers that drove the strongest halting phenotypes. By comparing available or generated stochastic labeling images (we performed this for VT012408-Gal4 as described in^26^, MCFO images were available for R36G02-Gal4 and R37F06-Gal4 ^27^), we identified that these three lines labeled BRK neurons. To validate this further, we devised a split-Gal4 screen with R36G02-Gal4DBD and candidate BRK targeting p65-ADs identified by using the Neuronbridge toolkit^27^. This screen (Extended Data Fig. 1d) helped generate split-Gal4 drivers for labeling all 6 BRK neurons (Fig 1) or subsets of BRK neurons (Extended Data Fig. 1d,e). We could further restrict expression to front leg specific BRK neurons (BRK-1-2) by using intersection with Scr-LexA^73^ (Extended Data Fig. 1d-f, Supplementary Table 1).

### Optogenetic activation in untethered animals

All the optogenetic activation experiments in untethered animals were performed using 6-9 days old female flies. The flies were loaded in behavioral arena as described in^26^ (44mm bowl shaped arena made of 1.5% agarose gel). The videos were recorded as in ^26^ using FLIR BlackFly-S Camera (FL3-U3-13Y3M-C) at a resolution of 1280 X 1024 at 30fps. The camera was fitted with an adjustable focus lens (LMVZ990-IR) and NIR bandpass filter (Midopt BP850) to allow IR imaging without artifacts from visible light. A custom designed LED panel^26^ provided backlit illumination with IR (850nm), green (530nm) or red (630nm) light. Both intensity and pulsing of each wavelength could be independently controlled and synchronized to the recording camera via TTL pulses generated using a custom Ardruino circuit. The bowl chamber arena was backlit with IR light (850nm) for video recording. We also provided continuous dim green light of intensity 0.1mW/mm^2^ during all optogenetic activation experiments to avoid jumping response in flies due to sudden bright light exposure. The green light level was adjusted so that it did not drive optogenetic stimulation even in a very sensitive reagent (MDN>CsChrimson). All experiments were performed at 25°C unless stated otherwise. Videos were tracked using FlyTracker software^74^ and data was analyzed in Matlab.

#### Activation in free walking flies

Experimental flies were loaded in above described setup and allowed to walk freely so that they could be assayed for optogenetics induced halting (Fig1 a-c and Fig2). The light stimulation protocol consisted of red light pulsed at 50Hz (5ms pulse width, average intensity at arena surface of 0.8mW/mm^2^) in a sequence of 50s OFF 10s ON, repeated 5 times. 10s OFF followed by 10s ON was considered as one trial during analysis. Distance/Rotation calculations were performed only for a period of 2s after stimulation onset to capture the strongest part of the activation phenotypes.

#### Activation in powdered flies

Flies were covered with powder (Reactive Yellow 86, Santa Cruz Biotechnology, Cat # sc-296260) to induce grooming^26,56^. Powdered flies loaded in bowl-shaped arenas described above were assayed for walking initiation with red light intensity of 1.0mW/mm^2^ pulsed at 100 Hz. The light stimulation protocol consisted of 50s OFF 10s ON sequence, repeated 5 times. For analysis, 10s OFF followed by 10s ON was considered as one trial.

#### Data Analysis

FlyTracker^74^ output was used for quantifying translational and angular velocities as in^26^. Angular velocity values correspond to “absolute angular velocity”. Rotation is defined as integral of angular velocity as in^26^. Distance and rotation was calculated for 2s period since stimulation onset. Pivots were defined as time periods with high angular velocity (>2r/s) and low translational velocity (<5mm/s) after smoothing with 0.5s window. These pivot thresholds are set based on empirical observation as well as previous literature^75–77^ indicating values for slow walking and high turning.

### Optogenetic silencing in untethered animals

All the optogenetic silencing experiments were performed using 6-9 days old female flies (unless stated) in the same setup and tracking/analysis pipeline as for activation experiments. The 530nm green LED used for silencing was adjusted to 0.65mW/mm^2^ average intensity at the arena walking surface to silence neurons expressing GtACR1.

#### Silencing in free-walking flies

Flies were loaded in bowl-chambers kept at 30°C (to elevate baseline walking) and assayed for decrease in walking velocity on silencing. The light stimulation protocol consisted of 60s OFF 30s ON sequence, repeated 3 times, for video duration of 5min.

#### Silencing in powdered intact flies

Flies were powdered as described above and assayed for interruption of grooming. The green light stimulation protocol consisted of 60s OFF followed by continuous ON for a duration for 6min.

#### Silencing in powdered decapitated flies (assay for tripping quantification)

The flies were decapitated using forceps and the neck was sealed using UV cured glue (Bondic). Flies that recovered well from this procedure (based on good self-righting and grooming behaviors) were chosen for experiments. The experimental flies were powdered (as mentioned above) and loaded in flat floor arenas (50mm diameter and 3mm height described in ^26^). The light stimulation protocol consisted of 30s OFF 10s ON sequence of green light (0.65mW/mm^2^), repeated 3 times, for video duration of 3min. Number of flies that trip during the light ON period were recorded.

#### Data Analysis

Velocities, distance and rotation quantification is as stated above. Stopping events were defined as instances where smoothed translational velocity (1.5mm/s) was below a threshold defined previously ^37^. Tripping events were quantified manually.

### Feeding related neuronal silencing assays

For inducing starvation, 1-3 days old female flies were transferred to retinal food and allowed to feed for 4 days. Flies were then wet-starved with 0.4mM retinal in water before testing.

#### Sugar preference assay

This assay was performed based on setup and conditions described previously^51^. 36 hours starved female flies were loaded in flat-floor circular behavioral arenas (50mm diameter) described above. The circular arena was covered with 2 halves of semi-circular filter paper, which were soaked with either water or 2M sucrose and left to dry overnight. Green light for GtACR1 based silencing was provided as described above. Flies were introduced in the chamber and allowed to explore and choose a preferred side for a duration of 4min. Video recording, tracking and analysis was performed as described above.

#### Sucrose-blob interaction assay

A 5µl drop (3mm diameter) with 1% agarose solution containing 200mM sucrose was placed in center of the flat circular arena (50mm diameter). One fly per arena was loaded and allowed to explore and find the sucrose drop. Green light as described above was provided throughout assay duration for GtACR1 based neuronal silencing. Flies that encounter the food before the video recording started were discarded from the analysis to ensure capturing the first feeding bout. Video recording and tracking was performed as described above. A fly within 3mm from the center of the food blob was considered as interacting with it (indeed, both the food blob and the fly are similar size i.e. ∼3mm). The food-zone was then defined as 6mm diameter circle centered around the food blob (Fig 5f). The exact frame corresponding to when the fly first found the food was manually annotated, and the data was aligned to this time point (Extended Data Fig. 8c). Quantification of food-zone stopping and velocities was performed within 5s of finding the food. Stops were defined as stated above.

#### Proboscis Extension Response (PER) Assay

This was performed as described in ^50^. Data was analyzed using Fisher’s Exact test between test and control genotypes.

### High-resolution 3D leg kinematics analysis

#### The setup and standard activation experiments on intact flies (Fig2)

The setup was based on previously described setups ^78–80^. Male flies aged between 7 to 10 days and starved for 6-8 hours were tethered to a 34-gauge needle with UV cured glue and placed on a spherical treadmill (6mm diameter) suspended in a stream of compressed air. The flies were placed on the ball with minimum wait time after tethering and allowed up to 5min of recovery before starting the experiment. The compressed air was passed through an in-line heating element to bring the local temperature on the ball up to 32°C to induce high speed spontaneous walking in the flies.

Each fly was left on the ball for a maximum of 20min during which time the forward, sideways and rotational velocity of the ball was monitored in real-time using two orthogonally placed motion sensors at 50Hz^78^. Each trial was triggered in a closed loop fashion when the forward velocity crossed a threshold empirically determined to signify sustained walking. Each fly could trigger a maximum of 10 trials, but flies that triggered more than 5 trials were included in the final dataset for analysis. Each trial was 7s long where the middle 3s had optogenetic stimulation using a red LED (625nm, Thorlabs M625F2) delivered to the fly via an optic fiber (Thorlabs FT400UMT). The LED was pulsed at 66Hz (2ms pulse-width) for all experiments, with a power of 0.23mW/mm^2^.

Eight cameras (FLIR BFS-U3-16S2M-CS) fitted with InfiniStix 194100 lenses and NIR bandpass filters (Midopt BP850) were placed surrounding the ball so that all legs were visible from at least a pair of cameras at all times. The fly was illuminated with a custom IR ring emitting focused light to the plane of the ball^78^. The cameras, IR light source and the ball tracker were all synchronized and triggered by an Arduino at 200Hz, with camera exposure time set to 200µs. The fly was recorded with a resolution of 1440×1072 pixels.

#### Camera Calibration, pose tracking and 3D position reconstruction

We chose 33 points of interest on the body of the fly to track using DeepLabCut (DLC) version 2.2.3 ^35^ – the notum, 2 wing hinges and 5 joints per leg (thorax-coxa, coxa-trochanter, femur-tibia, tibia-tarsus, and the tip of the tarsus). Separate ResNet-101 based neural networks were trained for all cameras (500,000 iterations), except the 3 front facing cameras which were all handled by the same network (5 networks in total). ∼830 manually annotated frames per camera were used for initial training of all networks (46 frames each from 18 flies) with a test-train split of 95-5%. An additional round of training was needed using ∼600 frames each (40 frames from 16 flies) before the tracking was satisfactory (error in pixels < 4 pixels for all networks).

The cameras were calibrated using the calibration module in the package Anipose^36^. We used a precision manufactured ChArUco board identical to the one used in Anipose publication as a calibration target^36^. The board was imaged at 15Hz from all cameras simultaneously at maximum resolution (1440 x 1072). When board-based calibration alone failed to give good results (as measured by the re-projection error in pixels of the final 3D model output of Anipose), animal-based calibration module was used to bring the reprojection errors below 20 pixels for all points. Anipose was used to triangulate all points, as well as calculate 8 angles per leg (Thorax coxa abduction, flexion & rotation, Coxa-Femur flexion & rotation, Femur-tibia flexion & rotation, Tibia-tarsus flexion).

#### Ball Fitting & swing-stance prediction

Using the tarsal tip positions time series from each fly, we arrived at an initial guess for the position of the ball center. The vector joining the ball center to the plane containing the thorax-coxa points was assumed to be normal to the plane, pointing in the opposite direction as the vector joining the plane to the notum. The magnitude of the vector was calculated as 4.5 times the distance between the wing hinge for each fly. Then this initial ball guess was optimized such that most points from the tarsal tip tracking are present on the surface of the ball with no points present inside the ball. The swing and stance phases were then estimated on the basis of the tarsal tip being on the surface of the ball or lifted away from it in any given frame. We estimated the step cycles based on the proximity of the tracked tarsal tip coordinates to the ball surface. We fit a sphere to the 3D reconstructed tarsal tip coordinates to estimate ball position. The position of the sphere in space and its radius was optimized iteratively using the squared distance of the tarsal tips to the surface of the sphere. Tarsal tip positions within 0.05% of the radius were considered stance, others swing. Furthermore, swing and stance phases less than 10ms, were filtered out.

#### Kinematic analysis

Only trials where the fly was walking with an average pre-stimulus velocity (in the initial 2s of the trial) greater than 1mm/s were considered for all leg kinematic analyses. The whole dataset was segmented into cases where the onset of optogenetic activation coincided with an ongoing swing phase or stance phase.

#### Definition of stopping bout

A period when the average ball velocity over 250ms was below 0.8mm/s.

#### Swing duration quantification

The pre-stimulus swing duration was calculated as the median swing duration from all swing events in the pre-stimulus period. The post stimulus swing duration was the duration of the swing that was in progress at stimulus onset.

#### Decapitated fly leg kinematics

The flies were decapitated as described above and tethered to the 34-gaguge needle with UV cured glue and placed on the ball.

#### MDN and MDN+BRK experiments

Every decapitated fly was subjected to 10 trials, each trial spanning 7s with optogenetic activation at 66Hz in the middle 3s, with a power of 0.23mW/mm^2^.

#### BDN2 and BDN2+BRK experiments

Every decapitated fly was subject to 4-5 different sets (10 trials each) with different intensity of red light (BDN2 expression levels varied and therefore 4 intensity levels were used per set: 0.013, 0.033, 0.15, 0.23, 0.34mW/mm^2^). Only sets where the flies showed walking initiation or resistance were retained for the co-activation set, in addition to a lower intensity where there was no visible movement.

### Functional connectivity

For functional connectivity experiments, whole CNS (brain+VNC) of female flies (6-9 days old) were dissected and imaged in extracellular saline (ECS) solution bubbled with carbogen^26^. Tissues were transferred on a PLL coated coverslip fixed in an imaging chamber (ALAMS-518SWPW) and imaged under a Bergamo II two-photon microscope using 20x NA= 1.0 objective lens (XLUMPLFLN, Olympus). During the entire imaging session, bubbled ECS with carbogen was delivered over the brains via a perfusion system (78018-40, Masterflex). GCaMP signal was recorded with a 920nm Ti:Sapphire laser (MaiTai DeepSee, Newport Spectra-Physics). For CsChrimson activation, a fiber-coupled 655nm LED (FC1-LED, Prizmatix) was positioned with a micromanipulator (Misumi XYZFG2) to deliver red light (0.04-0.26mW/mm^2^) onto the tissues. A given session typically consisted of 3-4 activation trials. The LED was synchronized with resonance imaging scanner (8.3kHz) such that it was only on during the non-imaging fly-back time of the scanner. This ensured that no light artifact appeared in the region of interest. A 2s stimulus was delivered at 50-100Hz, with an inter-trial interval >10s. Single-plane imaging was performed at a rate of 6Hz for BRK-BON1 experiment (field-of-view focused around BON-1 soma, Extended Data Fig. 6b-c), or 3Hz for Gr5a-FG/BB experiments (field-of-view focused around FG/BB neurites in the SEZ). Activation of the red-light LED was controlled and synchronized to the imaging system using ScanImage software (Vidrio Technologies). Background subtracted imaging data was analyzed using ImageJ and Matlab as in ^26^. ΔF/F = (F-F0)/F0, where F0 is pre-stimulus baseline (3s period before stimulus).

### Muscle imaging

To test the influence of BRK activation on leg muscle activity, we performed one-photon and two-photon calcium imaging of front leg femoral muscles. Female flies (5-8 days) of the genotypes BRK-Gal4>UAS-CsChrimson; MHC-LexA>LexAop-GCaMP6f^47^ were cold anesthetized and placed on a circular coverslip fixed in an imaging chamber (CSC-25L, Bioscience Tools). The wings and five legs – except the left front leg – were ablated. We applied UV-cured glue around the fly body and on the proboscis to avoid movements during imaging. The remaining front leg was glued either in its flexed (Fe-Ti angle∼ 10-27 degrees) or extended (Fe-Ti angle∼ 123-147 degrees) position. The fly holder was then flipped, such that the fly was underneath the coverslip, on opposite side to the objective. We could then add water and image with a water immersion objective (20x NA= 1.0 objective lens, Olympus XLUMPLFLN), while the fly remained dry on the other side of the coverslip. Muscle GCaMP signal in the front leg was recorded using both 1P wide-field fluorescence imaging (488nm mounted LED; Thorlabs) and 2P imaging (920nm Ti:Sapphire laser; MaiTai DeepSee, Newport Spectra-Physics). For activating CsChrimson expressed in BRK, a 655nm LED (FC1-LED, Prizmatix) was placed with a micromanipulator and used to deliver red light (0.07mW/mm2) towards the fly thorax. A given session typically consisted of four activation trials for both one- and two-photon experiments.

In one-photon experiments, red light was delivered for 2s with 10s inter-stimulation intervals. Muscle GCaMP signal was acquired at a frame rate of 50Hz. Red light stimulation was controlled via TTL inputs synchronized to the imaging session using ThorCam software (Thorlabs) plus an external Arduino based trigger box (Thorlabs TSI-IOBOB2). In two-photon experiments, the red light stimulation (100Hz) was as described in functional connectivity experiments. Muscle GCaMP signal was acquired at 6Hz.

For tibia-movement, we used protocol described in^48^. Briefly, a fly was mounted on a coverslip as described above. A magnetic pin (1mm length) was then glued on the front left leg tibia. A magnet mounted on a programmable servo motor (Silver max Hybrid Servo Motor, Precise Motion and Control Inc.) was used to forcibly flex and extend the femur-tibia joint (1s per flexion/extension, repeated 4 times). Femoral muscle activity was imaged under epifluorescence (1P) while delivering optogenetic stimulation using the 655nm LED described above, during the entire session. Muscle GCaMP signal was acquired at a frame rate of 50Hz. A simultaneous video of the tibia movements (720×540 resolution; 50Hz) was acquired in SpinView software (FLIR) and synchronized to the imaging session using ThorCam software (Thorlabs) plus an external Arduino based trigger box (Thorlabs TSI-IOBOB2).

For optogenetic stimulation experiments, data were analyzed as for functional connectivity. For tibia movement experiments, data were analyzed similarly except in this case F0 was pre-movement baseline (median of 15 frames (-20 to -5) before movement initiation).

### *In-vivo* Imaging

Female flies (4-7 days) of the genotypes BRK-Gal4>Act88F:Rpr; UAS-CG6f-tdTomato or BDN2-Gal4>Act88F:Rpr; UAS-CG6f-tdTomato were anesthetized on ice and tethered on a custom fly holder^26^ Thoracic dissection for VNC imaging was then performed as described in ^17^. Artificial Hemolymph (AHL^81^) solution was used during dissection and imaging of the exposed VNC. After dissection the holder was placed under a Bergamo II two-photon microscope (Thorlabs) under a water immersion objective (40x NA= 0.8 objective lens Nikon CFI APO NIR), and an air-supported ball was positioned under the fly in a similar setup as described above for leg-kinematics analysis. Flies with uncoordinated leg movements (∼25%) were discarded before the experiment. After an acclimation period to the ball (∼15-20min), a volume containing axonal projections from BRK or BDN2 was imaged using at a volumetric rate of 2Hz (BRK) or 6Hz (BDN2), using 920nm Ti:Sapphire laser and fast z-piezo device. Video of the behaving fly (720×540 resolution; 200Hz) was acquired in SpinView software (FLIR) and synchronized to the imaging session using ScanImage software (Vidrio Technologies). The synchronized calcium imaging and ball velocity data were analyzed offline. For BRK imaging experiments, additional behaviors were manually annotated (Fig. 6e-h) using FlyTracker software^74^. Data analysis was performed using custom scripts in python and Matlab.

### Immunohistochemistry

All CNS dissections and immunohistochemistry were performed as described in ^82^ with detailed protocols available at https://www.janelia.org/project-team/flylight/protocols. Primary antibodies used were chicken anti-GFP (1:1000, Thermo Fisher Scientific, AB_2534023), rabbit anti-dsRed (1:500, CloneTech, AB_10013483) and anti-Bruchpilot (1:500, nc82, mouse monoclonal, Developmental Studies Hybridoma Bank, AB_2314866). Alexa fluor secondary antibodies (Thermo Fisher Scientific) were used at 1:500 dilution (Goat anti-chicken: Alexa488, AB_2576217, Goat anti-rabbit: Alex568, AB_10563566, Goat anti-mouse: Alex568, AB_2534072 and Goat anti-mouse: Alex647, AB_141725).

### Fluorescent In-Situ Hybridization (FISH)

This was performed as a part of large scale FISH imaging by Alyson Petruncio and FlyLight team at Janelia Research Campus, as described in ^42^.

### Connectome constrained modeling

The neuronal activity was simulated as a spiking neural network in the brian2 software v2.5.1^83^. Details about the model, including the original code, are described elsewhere^4^. The original code has been modified to allow for stimulating and silencing neurons at arbitrary time points throughout the simulation. Briefly, a leaky integrate-and-fire model is constructed based on the fully annotated connectome^5^. The connection strength between two neurons is proportional to the number of synapses linking them. Neurotransmitter predictions based on the EM dataset^41^ determine whether the interaction between two neurons is inhibitory (GABA, Glu) or excitatory (all other). Neuronal stimulation can be mimicked in the model by adding a Poisson spike train of defined frequency as an input to a neuron. Neuronal silencing can be mimicked by simply severing all outgoing synaptic connections of the desired neuron. It is important to realize that the intrinsic firing rate of neurons in this model is zero and that the Poisson inputs constitute the only external input. The firing rates reported here are averaged over 30 trials, 1000ms each with 0.05ms integration time steps.

Related to Fig 3: The walk neurons P9/BPN were stimulated bilaterally at 150Hz/50Hz, respectively. The firing rates for the top 100 neurons are shown in panel 3a/3e. Panels 3b-d/3f-h show the same neurons as 3a/3e, but here the stop neurons BB, FG, and BRK, respectively, are stimulated at 150Hz between 0.25 and 0.75s. All rates in the top panels 3a-h were smoothed with Gaussian kernel (sigma= 25ms). In the wiring diagram shown in bottom panels 3a-h, only the most active DNs with firing rates >10.3Hz/>20Hz for P9/BPN stimulations are shown. The node color shows the average firing rates of the most active DN during walk neuron stimulation (Fig. 3a,e) and during walk-halt co-stimulation (Fig. 3b-d and Fig. 3f-h) normalized to the maximum firing rate in 3a and e, respectively. As BPN projects contra-laterally, the left hemisphere DNs receiving input from contralateral BPNs (right) were chosen for representation (this applies to Extended Fig 10c,d, where walk neurons BPNs and P9 are stimulated at 150 and 50Hz, respectively, and oviDN and MAN1 are stimulated at 150Hz).

Related to Extended Data Fig 6: g,h shows all neurons that are differentially affected in walk neuron stimulation and walk-stop neuron co-stimulation. Walk neurons BPN and P9 are stimulated at 150Hz and 50Hz, respectively, stop neurons FG and BB are stimulated at 150Hz.

Related to Fig 5: The GRNs were activated (150Hz) in Fig. 5a and downstream feeding pathway with increased firing rate along with FG and BB are shown as wiring diagram. The node colors correspond to the firing rate except for oDN1 and BDN1 (for simplicity, the graphs are limited to one hemisphere). The heatmaps in panel 5c and 5d show the average firing rate of oDN1 and BDN2 normalized to the maximum value in the left panel, respectively. Left: co-stimulation of left hemisphere sugar GRNs and bilateral P9 or BPN walk neurons. Middle: same as left, but while silencing stop neuron FG bilaterally. Right: difference in firing rate between left and middle panels.

Cytoscape^84^ was used for creating all the wiring diagrams shown in this study. The edges in the diagram illustrate the connectivity in the model (red: excitatory, blue: inhibitory). The arrow size represents the connection strength, (the connections below 5 synapses are excluded). We show only one hemisphere for simplicity except in case of Extended Data Fig. 2a and Extended Data Fig. 6a. The node colors represent firing rates wherever mention.

#### Statistics

All statistical tests were performed in Matlab or Graphpad Prism. All 2-group comparisons (unless indicated otherwise) were performed using non-parametric Mann-Whitney test. All multiple-group comparisons were performed using non-parametric Kruskal-Wallis test followed by Dunn’s multiple comparison with appropriate control. Exact sample size values for each plot are reported in Supplementary Table 1.

#### Code

Custom scripts were written in either Matlab 2021a or Python. A Github repository containing all code related to this manuscript will be made available post-publication on the bidaye-lab Github page (pre-publication upon reasonable request).

## Author Contributions

Neha Sapkal performed all free-walking behavior experiments, analyzed modeling data and coordinated other experimental efforts with support from Kazuma Murakami, Gianna Vitelli and Benjamin Bargeron. Nino Mancini performed all functional imaging experiments with support from Kazuma Murakami and Nico Spiller. Divya Kumar performed all leg kinematics experiments and analysis with support from Kate Maier and Nico Spiller. Nico Spiller wrote the computational model under supervision of Philip Shiu. Katharina Eichler and Gregory Jefferis led connectomics efforts pertaining to all descending and ascending neurons in the brain and nerve-cord connectome data. Gabriella Sterne provided the SEZ-splitGal4 lines, Gr5a reagents and one functional imaging dataset pertaining to Gr5a activation. Salil Bidaye conceived the project, designed and supervised all the experiments, and wrote the manuscript with support from co-authors.

## Acknowledgements

We thank Kristin Scott for guidance during early phase of this project. We thank Ansgar Büschges, Silvia Arber and Vivek Jayaraman for providing critical feedback on this work. We thank Lesley Colgan for feedback on the manuscript. We thank Moritz Haustein, Till Bockemühl and Michael Dübbert from the Büschges lab for technical assistance with leg-kinematics analysis setup and pipeline. We thank MPFI mechanical workshop for custom designed parts in all our setups. We thank John Tuthill and Wei-Chung Allen Lee for providing early access to nerve cord connectome data (FANC). We thank Amy Sterling and Jasper Phelps for coordinating FlyWire and FANC consortiums and technical assistance. We thank Pavan Ramdya for sharing reagents for in-vivo nerve-cord imaging. We thank Alyson Petruncio and FlyLight team at Janelia Research Campus for FISH images. We thank Rod Murphy and Juan Lopez for performing preliminary muscle recordings and discussions. We thank the Princeton FlyWire team and members of the Murthy and Seung labs, as well as members of the Allen Institute for Brain Science, for development and maintenance of FlyWire (supported by BRAIN Initiative grants MH117815 and NS126935 to Murthy and Seung). We also acknowledge members of the Princeton FlyWire team and the FlyWire consortium for neuron proofreading and annotation. We acknowledge FlyWire proofreading efforts by Varun Sane, Laia Serratosa Capdevila, Alex Javier, Siqi Fang, Griffin Badalamente, Markus Pleijzier, Kai Feng, Quinn Vanderbeck, Ryan Willie, Doug Bland, Austin T Burke, Megan Wang, Regine Salem, Zairene Lenizo, Michelle Pantujan, Nash Hadjerol, J. Anthony Ocho, Miguel Albero, Darrel Jay Akiatan, Kendrick Joules Vinson, Mendell Lopez, Chitra Nair, Zeba Vohra, Bhargavi Parmar, Kaushik Parmar, Dhara Kakadiya, Itisha Joshi, Dhwani Patel, Dharini Sapkal, Arti Yadav, Anjali Pandey and Wolf Huetteroth. NM is funded by DFG Grant, Walter Benjamin Program, MA10161/1-1. Nico Spiller is funded by the Carl Angus DeSantis Foundation. Additional proofreading and infrastructure was supported by Wellcome awards 203261/Z/16/Z and 220343/Z/20/Z to Jefferis. SSB is funded by the Max Planck Society.

---

**Extended Data Fig 1:**
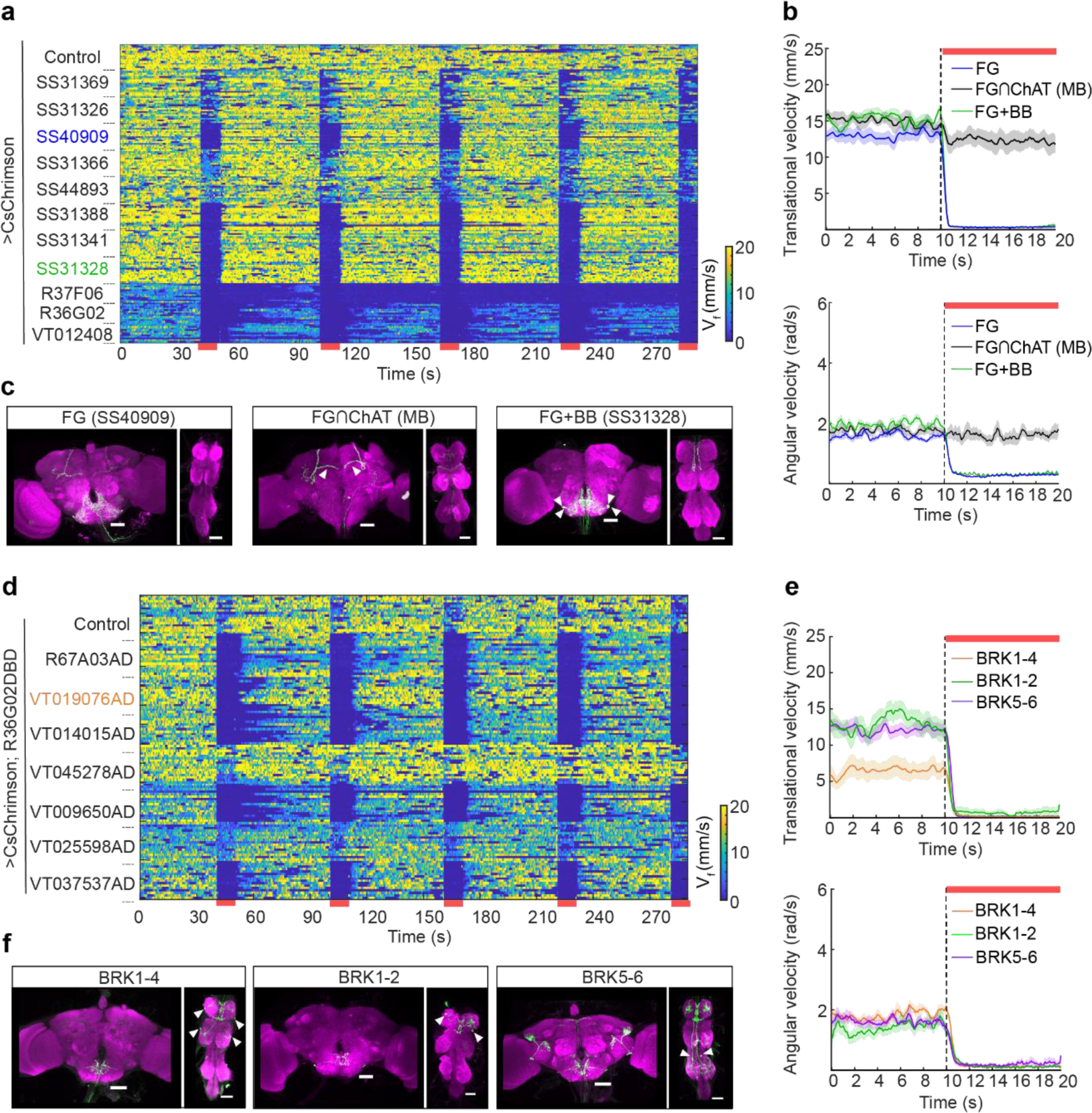
Neural activation screen to identify halting neurons: **a.** Translation velocity heat map of free-walking control flies and broad lines expressing CsChrimson. Red bars at bottom of heatmap indicate 5 trials of 10s red light driven optogenetic stimulation. n= 16 flies per genotype. **b.** Trial averaged translational velocity (top) and angular velocity (bottom) of FG (SS40909), FG∩ChAT (MB), FG+BB (SS31328). **c.** Immunohistochemistry images of genetic reagents used in (b); white arrowheads show mushroom bodies (MB) (CsChrimson-mVenus: green, neuropil: magenta, scale-bars: 50μm). **(d)** Translation velocity heat map (every row shows velocity of single fly) of free-walking flies with a focus on sparse genetic lines labelling BRK. Red bars at bottom of heatmap indicate five trials of 10s red light driven optogenetic stimulation. **(e)** Trial averaged translational velocity (top) and angular velocity (bottom) of free-walking flies with CsChrimson expressed only in BRK1-4, BRK1-2, or BRK5-6. 10s optogenetic stimulation (red bars) starts at vertical stippled lines in velocity plots. n= 10-16 flies per genotype. Table S1 shows full experimental genotypes. **(f)** Immunohistochemistry images of genetic drivers used in (e) (CsChrimson-mVenus: green, neuropil: magenta, scale-bars: 50μm). Supplementary Table 1 shows full experimental genotypes and sample sizes.

**Extended Data Fig 2:**
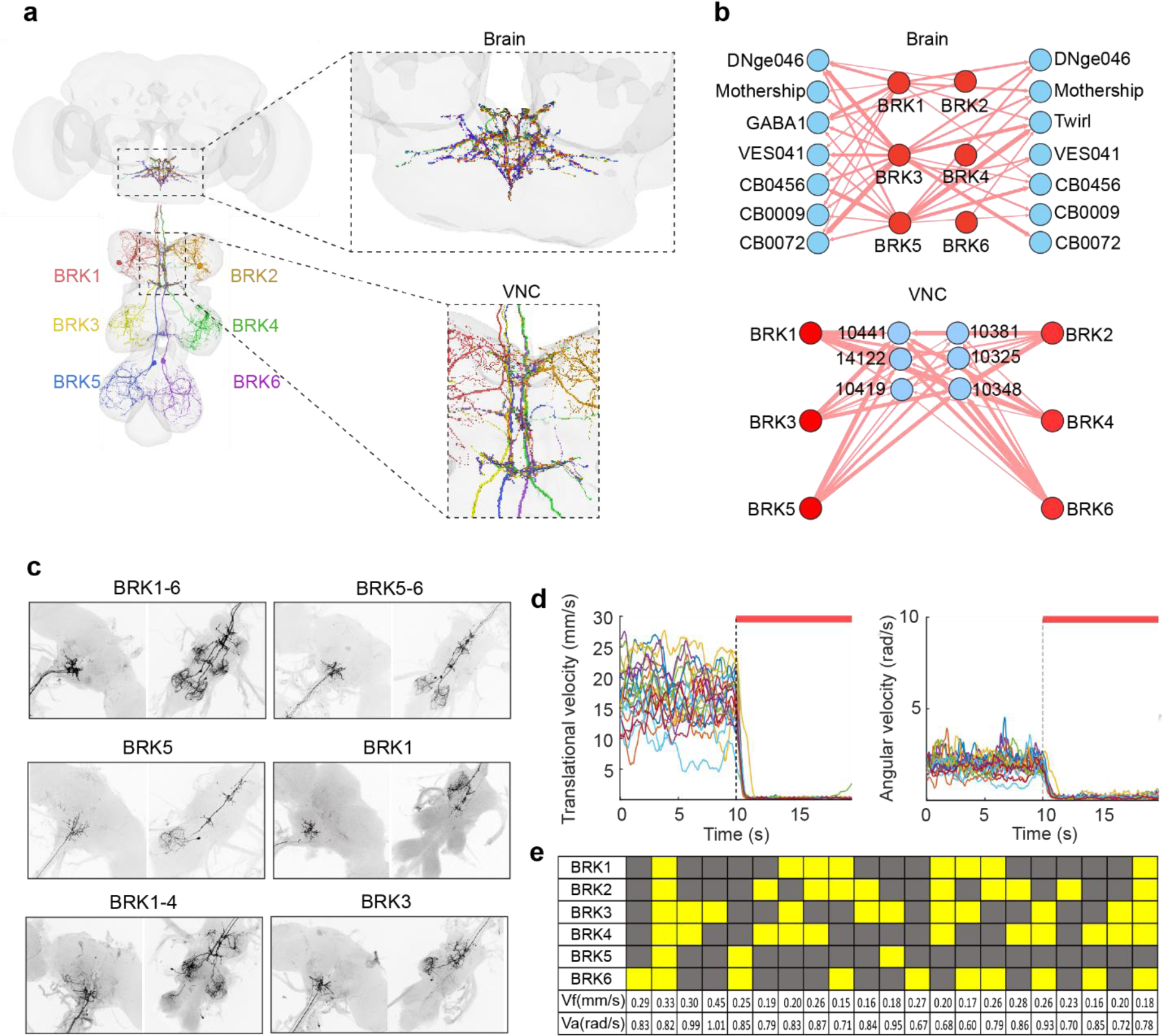
Segment specific BRK activation causes halting. **a.** EM segmentation of all six BRK neurons. Insets show BRK axonal projections in the brain (top) and VNC (bottom). **b.** Connectome based wiring diagram showing BRK major post-synaptic partners in the brain and VNC. Arrow thickness indicates synaptic strength (5-100 synapses for top and 5-400 synapses for bottom); red indicates predicted excitatory connections. **c.** Stochastic expression in distinct subsets of BRK neurons. **d.** Translational angular (left) and angular velocity (right) of individual, free-walking flies with different subsets of BRK optogenetically activated (red bars) using CsChrimson. 10s light stimulation starts at vertical line in velocity plots. n= 21 flies per genotype. **(e)** Summary table from data in (d). Each column represents expression in 1 fly (yellow: present, grey: absent). Supplementary Table 1 shows full genotypes and sample sizes.

**Extended Data Fig 3:**
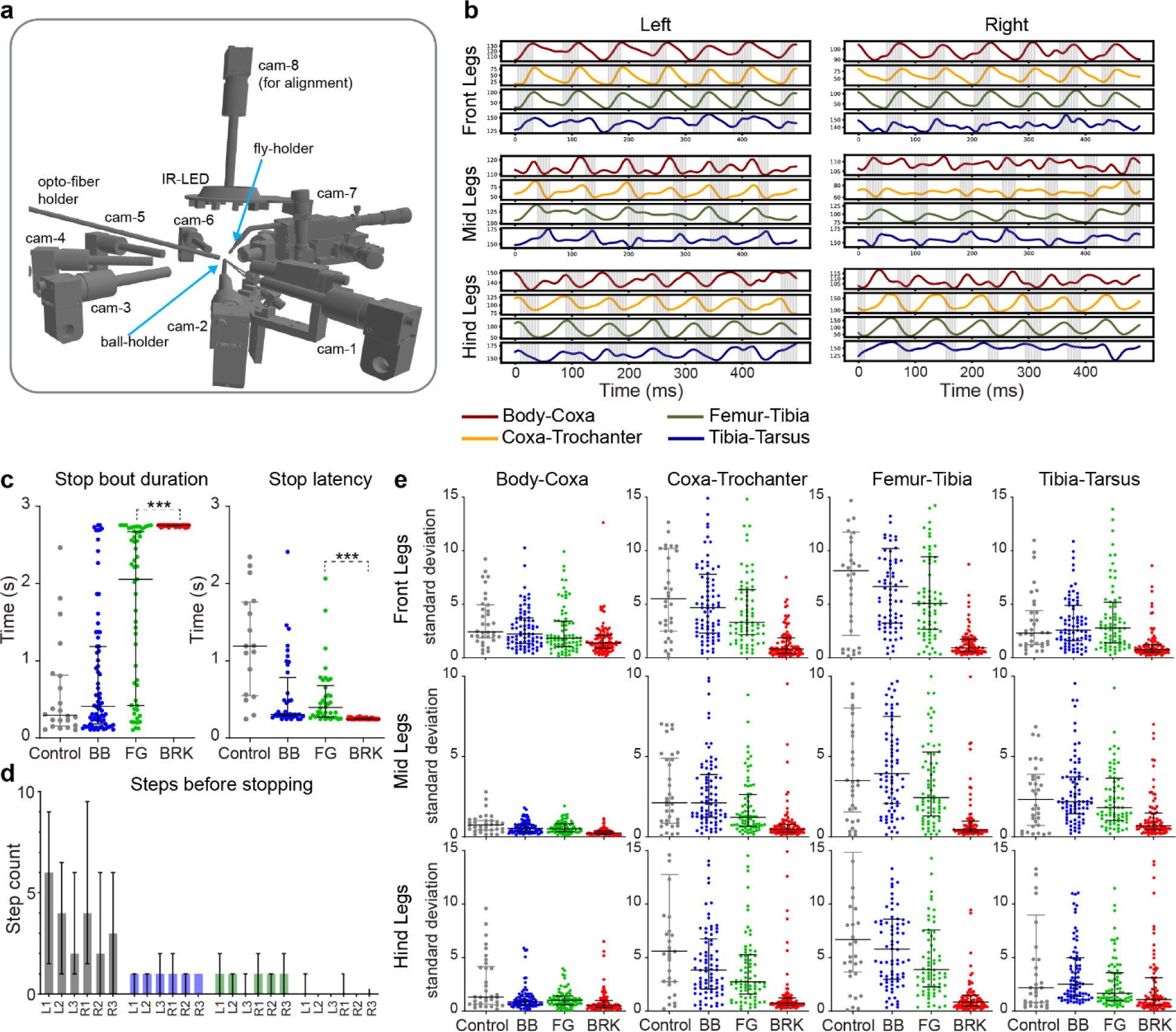
Leg Kinematics Analysis. **a.** Pipeline for high-resolution 3D leg kinematics analysis. The fly is tethered onto a custom made fly-holder and mounted on to a ball placed on the ball holder at the center. The fly is illuminated using a custom made IR-LED ring from above. Seven cameras image the fly from different sides and the top-down camera is used to align the fly on the ball. Two orthogonally placed motion sensors track the rotations of the ball allowing to calculate the ball velocity. Red light for optogenetic activation is supplied through an optical fiber mounted on the holder. **b.** Example traces of the four flexion angles (body-coxa, coxa-trochanter, femur-tibia and tibia-tarsus) calculated per leg, for all six legs in tethered walking flies. Grey shading corresponds to swing phase. **c.** Stopping bout durations (left) and stopping latency (right) during optogenetic activation of BB, FG and BRK. Mann-Whitney Test to compare between the strong halting phenotypes FG and BRK (n is in Supp. Table 1, *** p <0.001). **d.** Number of steps before flies stop upon optogenetic activation of BB, FG and BRK, compared to control flies. **e.** Standard deviation of all flexion angles calculated over the whole light-ON period, for the respective genotypes. Supplementary Table 1 shows full genotypes and sample sizes.

**Extended Data Fig 4:**
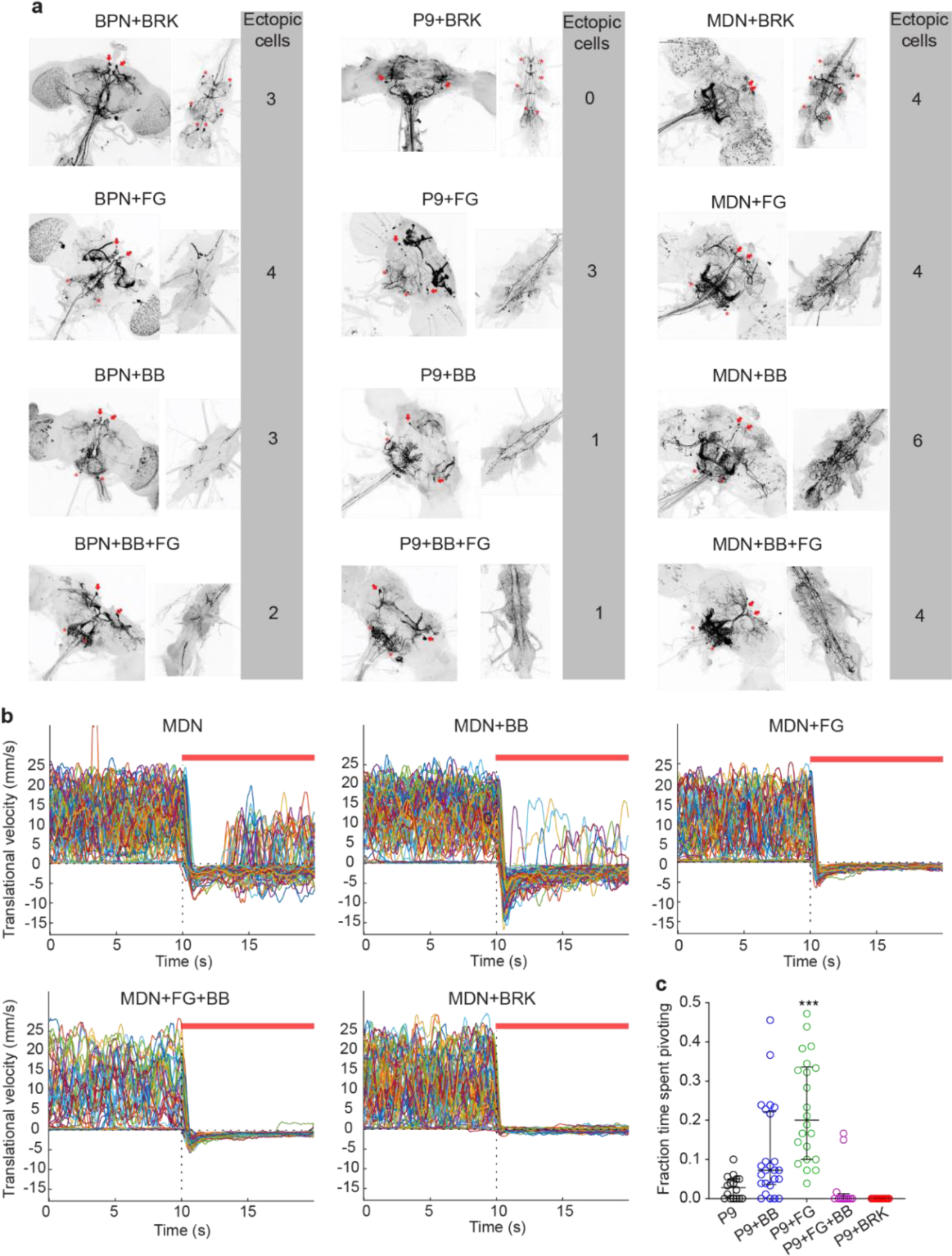
Walk-Halt Co-activation. **a.** Light microscopy images of combined split-Gal4 lines comprising walk neurons (P9, BPN, MDN) and halt neurons (BRK, FG, BB, and the FG+BB line). Arrows indicate walk neurons and asterisks show halt neurons. Ectopic expression is indicated as cell numbers. **b.** Per-trial translational velocity of pooled across all flies (n = 10-16 flies, 4 trials per fly) with MDN co-activated with BB, FG, BB+FG, or BRK. 10s light stimulation (red bars) starts at vertical stippled line in velocity plots. **c.** Fraction of time free-walking flies spent pivoting of upon P9 co-activation with BB, FG, FG+BB, or BRK. Dots indicate individual flies; n= 12-25, median±interquartile range. Kruskal-Wallis test followed by Dunn’s multiple comparison with genetic-background control (*** p < 0.01). Supplementary Table 1 shows full genotypes and sample sizes.

**Extended Data Fig 5:**
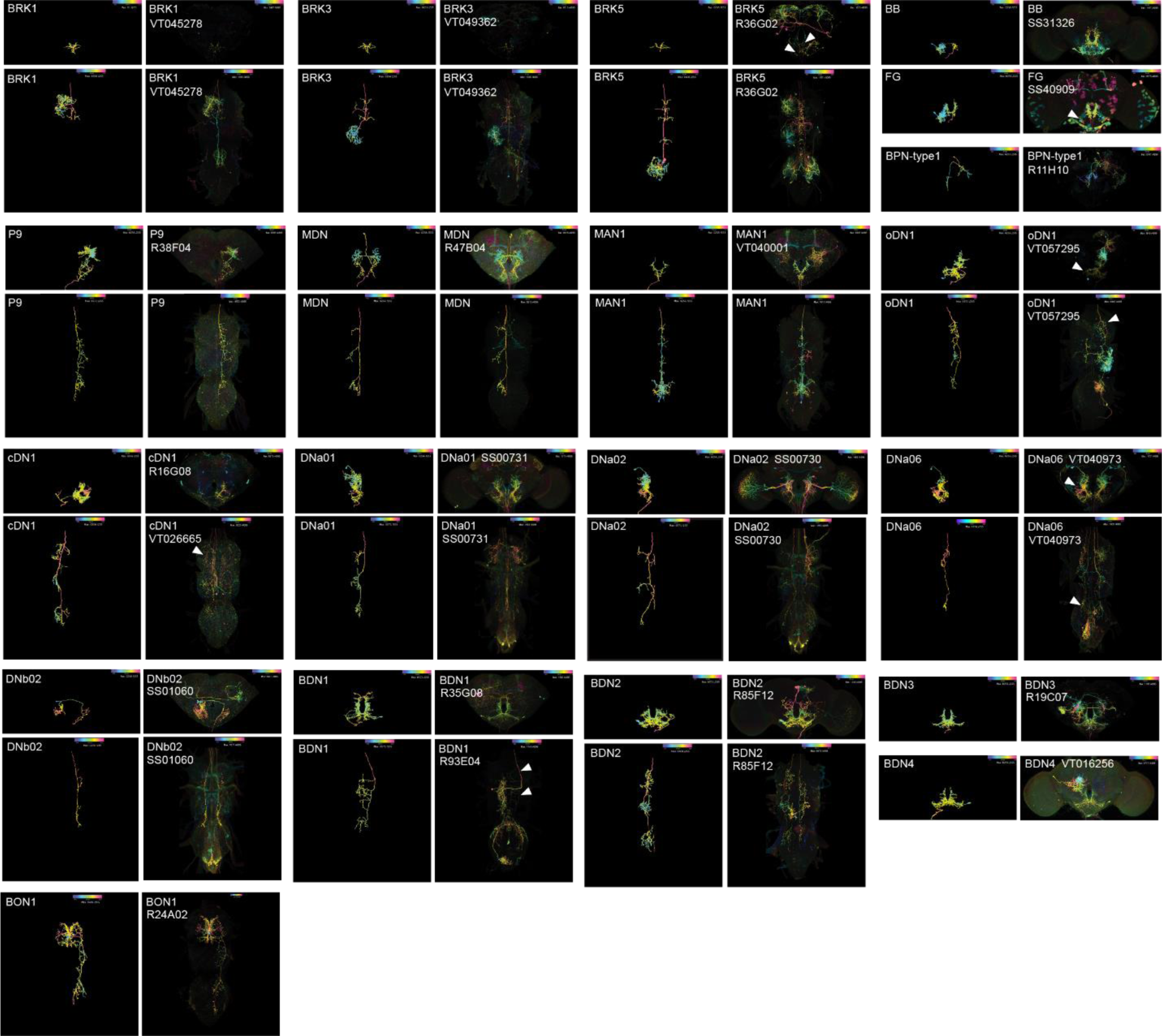
MCFO-colormip comparison (brain and VNC) for neurons of major interest in this study, grouped by halt and walk neuronal types, and by their major downstream synaptic partners. Nomenclatures to annotations given in light- and electron-microscopy datasets. White arrowheads show neuron of interest.

**Extended Data Fig 6:**
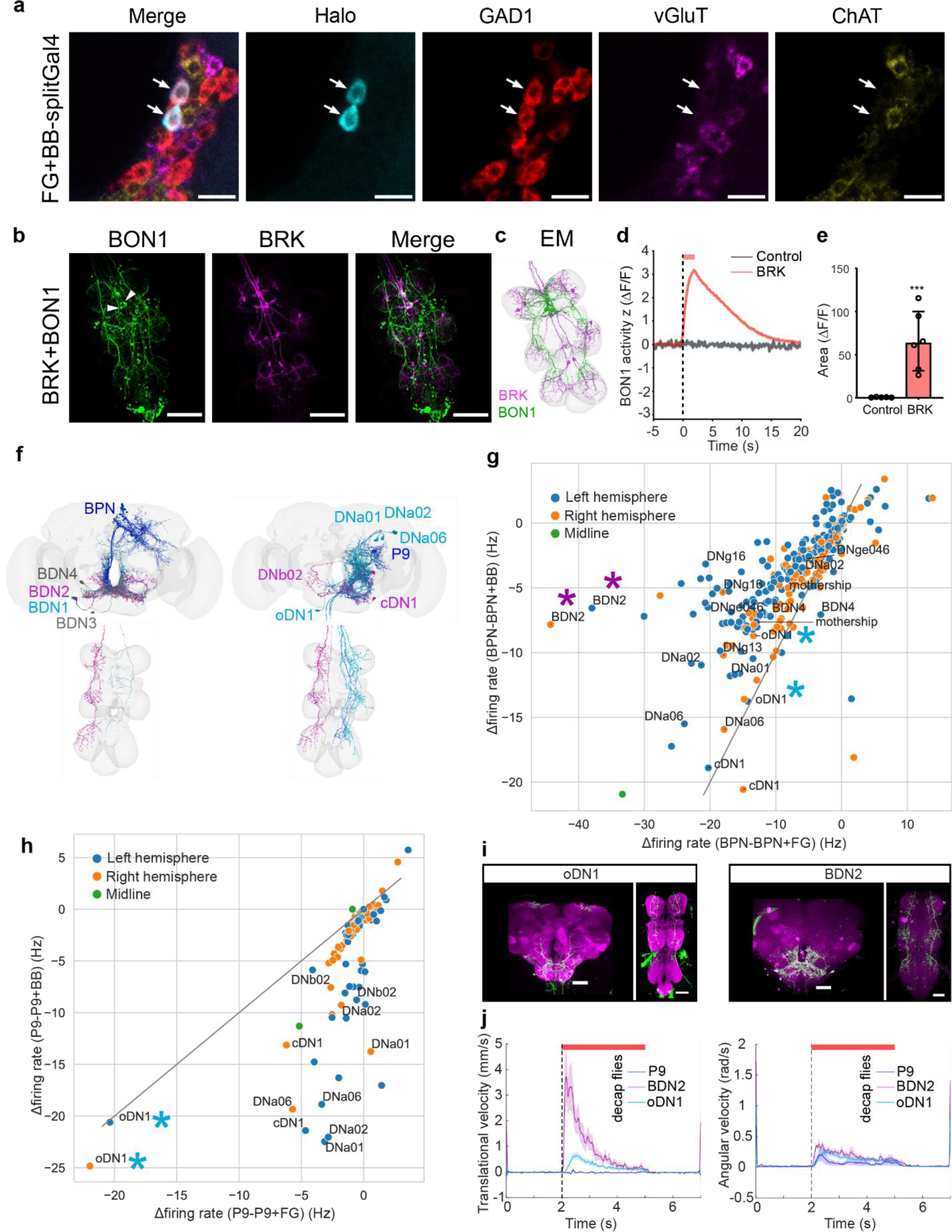
FG and BB inhibit specific nodes of BPN and P9 pathway. **a.** Neurotransmitter profile for the combined FG+BB split-Gal4 using FISH. Arrows indicate BB/FG soma. **b.** Light microscopy images of genetic line co-labelling BRK and BON1; (GCaMP6s: green, ChrimsonR-mCherry: magenta, scale bars: 100μm, white arrowheads indicate BON1 soma). **c.** EM segmentation of BRK and SON1 in the VNC. **d.** BON1 activity (GCaMP6s) in explant brains without or with BRK activated (ChrimsonR). 2s optogenetic activation (red bar) starts at vertical stipple line; n=5-6 flies per group. **e.** Area under the curve (ΔF/F) for control and BRK activated groups in (d). Mann-Whitney Test (*** p< 0.001) **f.** DNs recruited by BPN and P9 (magenta: contralateral descending; blue: ipsilateral descending; grey: unidentified descending arbors). **g,h.** Scatter plot showing DNs downstream of **(g)** P9 and **(h)** BPN differentially affected by FG versus BB. **i.** Expression pattern of split-Gal4 lines covering oDN1 and BDN2. (CsChrimson-mVenus: green, neuropil: magenta, scale bars: 50 μm). **j.** Trial averaged translational velocity (left) and angular velocity (right) of decapitated tethered flies, while activating P9, oDN1, or BDN2. 3s light stimulation (red bars) starts at vertical stippled lines in velocity plots. n= 7-10 flies per genotype, mean±SEM. Supplementary Table 1 shows full genotypes and sample sizes.

**Extended Data Fig. 7:**
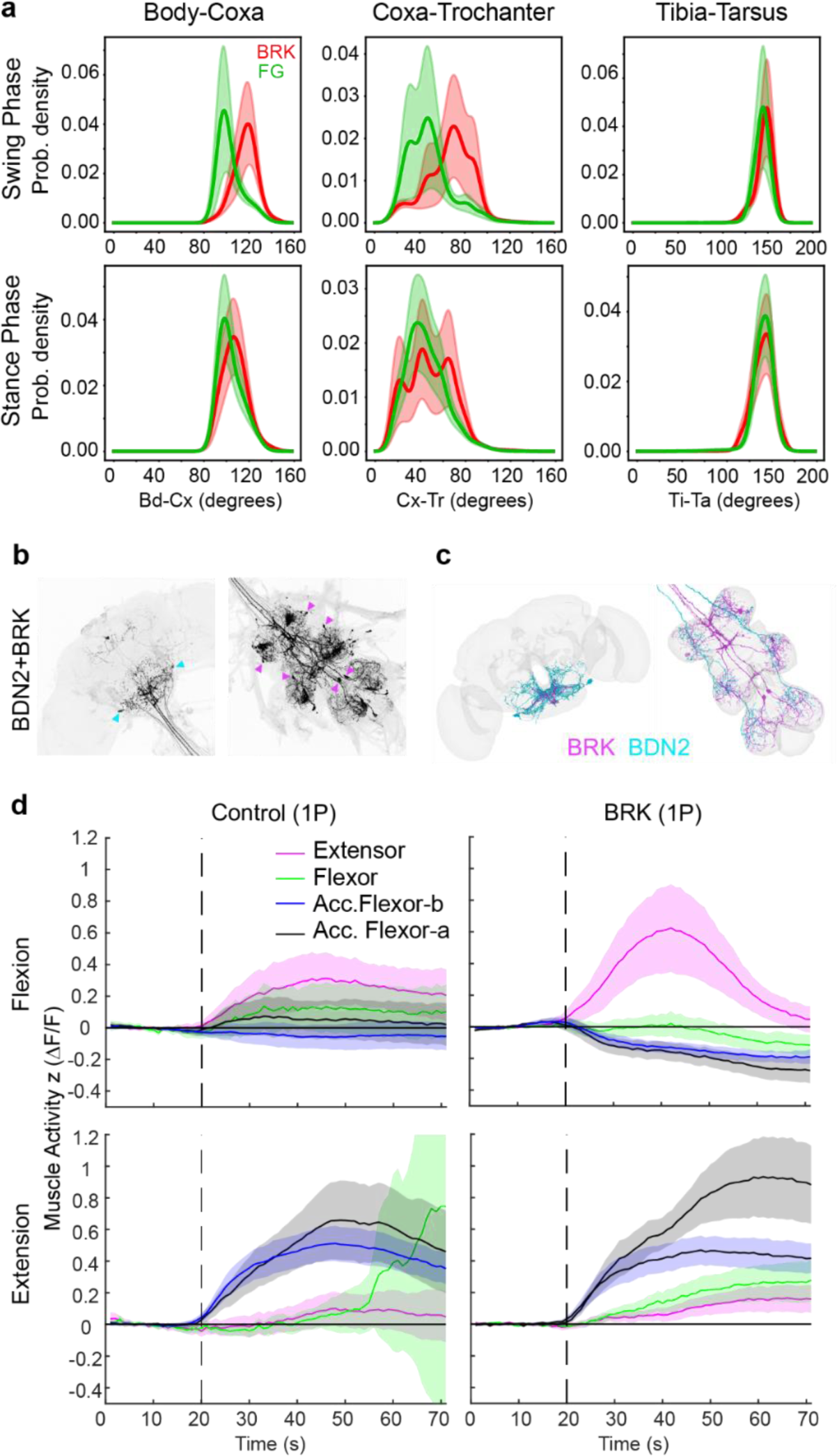
Brake Mechanism Analysis: **a.** Probability density of front leg joint angle (body-coxa, coxa-trochanter and tibia-tarsus) if BRK (red) or FG (green) was optogenetically activated during swing phase (top) or during stance phase (bottom). **b.** Light microscopy images of combined split-Gal4 lines for co-labelling BRK and BDN2. Colored arrowheads indicate soma of the corresponding neurons in the brain (left) and VNC (right). **c.** EM segmentation of BRK and BDN2 in the brain (left) and VNC (right). **d.** Left front leg femoral muscle activity (MHC>GCaMP6s, color coding as in Figure 4) while Fe-Ti joint is forcibly flexed (top) or extended (bottom) in control (left) or BRK>CsChrimson (right) flies during light stimulation (red bar) measured under epifluorescence (1P). 1s flexion/extension start at vertical stippled lines in plots. n= 5 flies per genotype. Supplementary Table 1 shows full genotypes and sample sizes.

**Extended Data Fig. 8:**
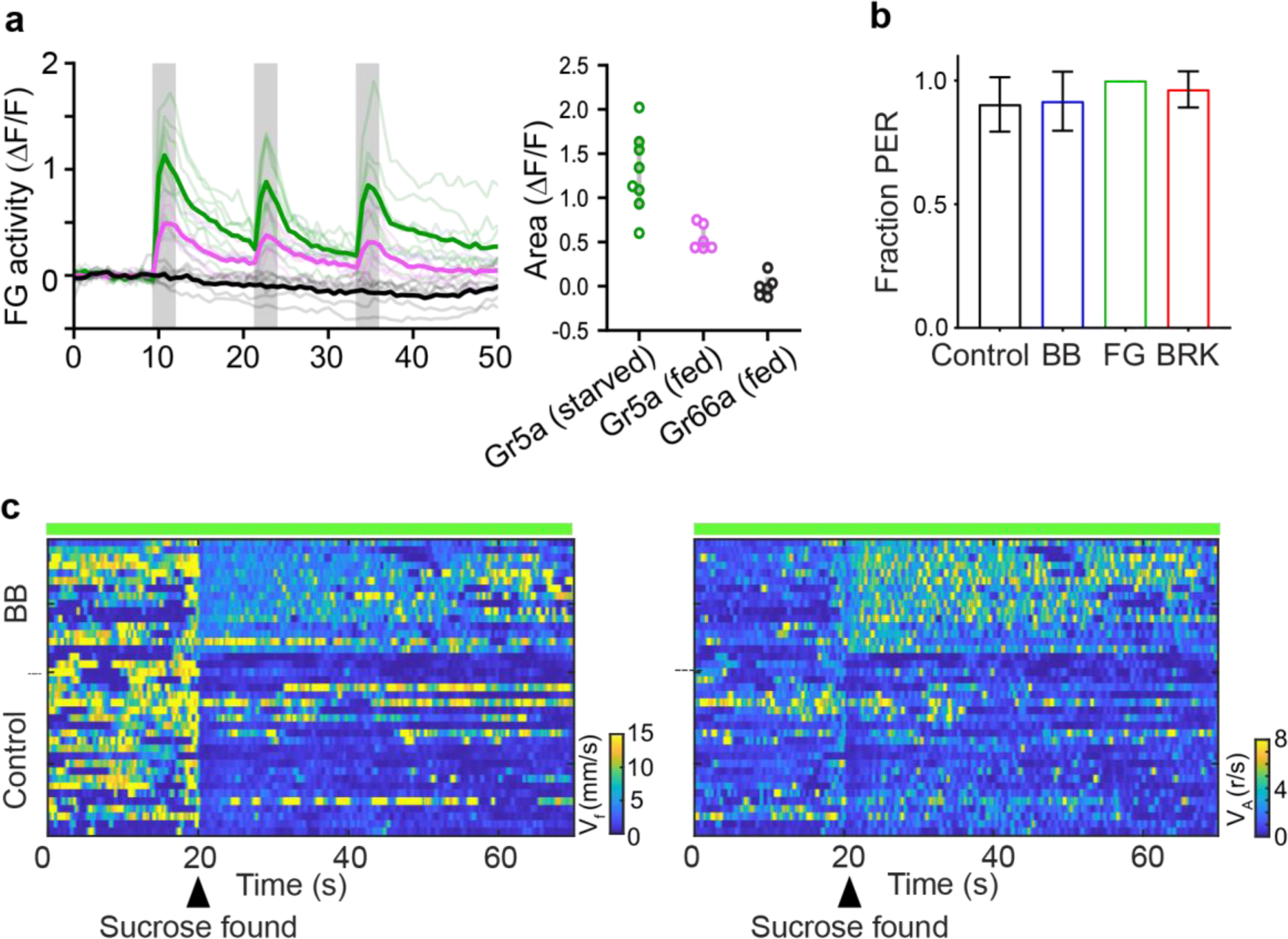
**a.** FG activity (ΔF/F, GCaMP7b) during optogenetic stimulation of sweet gustatory sensory neurons (Gr5a, in fed or starved flies) or bitter gustatory sensory neurons (Gr66a, in fed flies). **b.** Proboscis extension response (PER) upon sucrose presentations in flies with FG, BB, or BRK silenced (GtACR1) throughout, compared to control flies. n= 26-34 flies, Fisher’s exact test with control showed no difference for any genotype. **c.** Translational velocity (left) and angular velocity (right) heatmaps of starved, free-walking flies with BB silenced throughout (GtACR1), compared to control flies. Green bar on top of heatmap indicates green light stimulation. Arrow indicated time point where the sucrose was found by the flies. Supplementary Table 1 shows full genotypes and sample sizes.

**Extended Data Fig 9:**
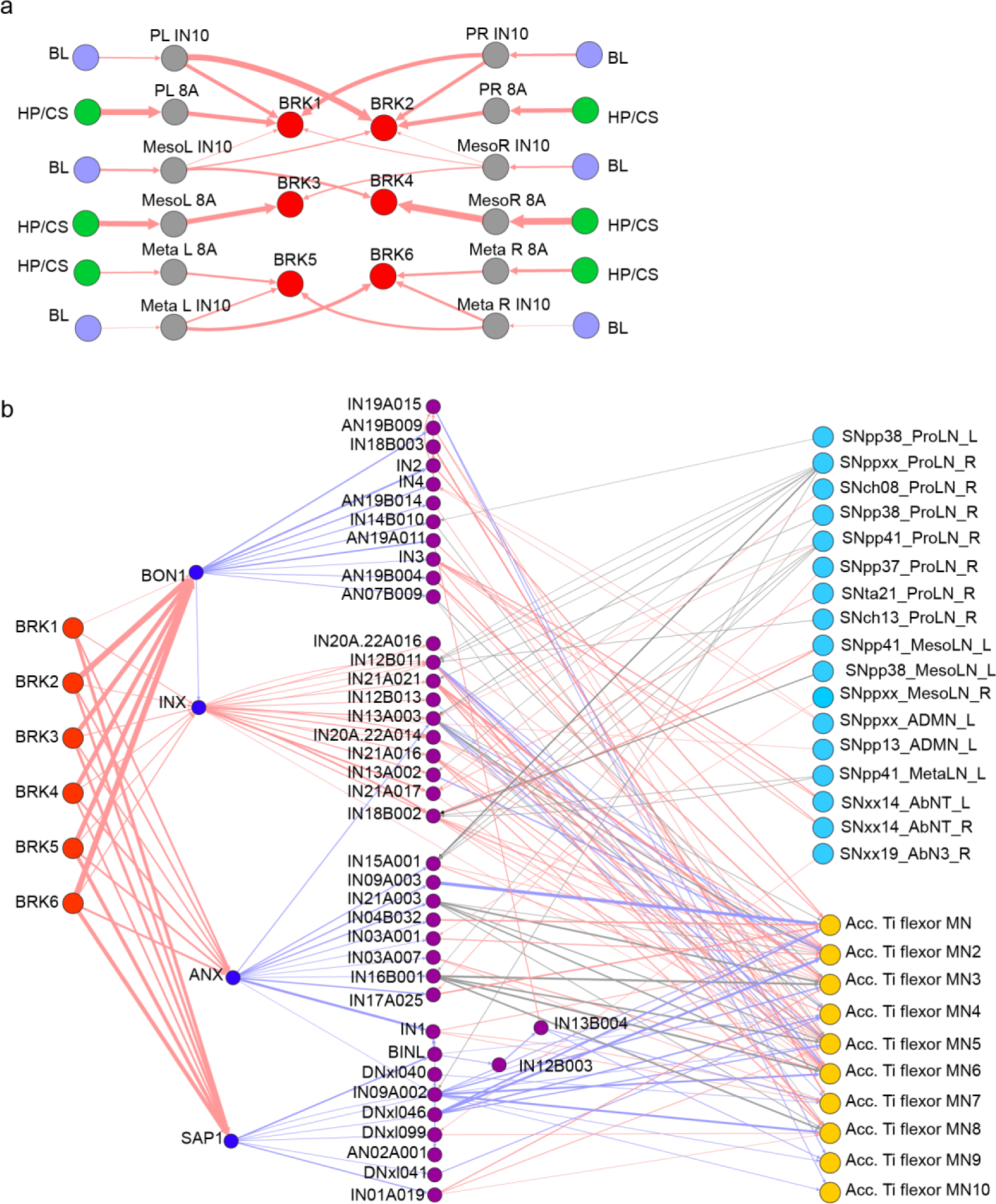
Connectome based analysis of BRK connectivity in the VNC. **a.** All 6 BRK neurons receive segment specific leg mechanosensory (BL: Bristle) and proporiosensory (HP: hairplate, CS: Campaniform sensilla) inputs. **(b)** 2^nd^ order downstream BRK output receive input from sensory neurons (mostly proporioceptive, blue) and provide output to accessory tibia flexor motoneurons (Acc. Ti flexor MN, yellow). In **a-b** arrow thickness indicates synaptic strength (5-300 synapses). Red indicates excitatory connections; blue indicates predicted inhibitory connections.

**Extended Data Fig 10:**
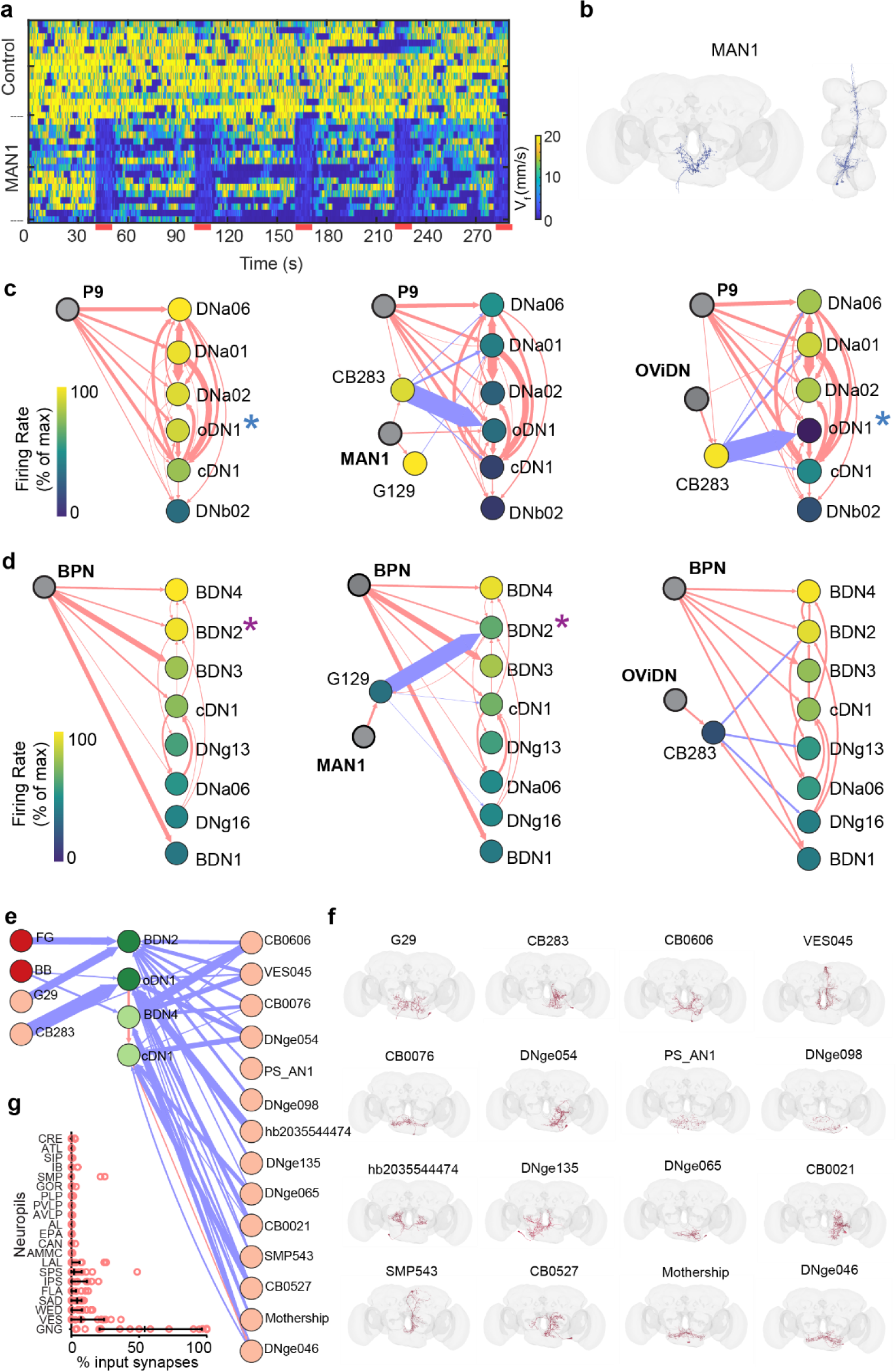
Prediction of novel halting pathways. **a.** Translation velocity heat map of free-walking control flies and flies expressing CsChrimson in MAN1. Red bars at bottom of heatmap indicate 5 trials of 10s red light stimulation. **b.** EM segmentation of MAN1 in the brain (left) and VNC (right). **c,d.** Simulation results for activating **(c)** P9, P9+MAN1, P9+OViDN and **(d)** BPN, BPN+MAN1, BPN+OViDN. Nodes in the wiring diagram are color coded based on firing rate in the full simulation, identical to the heatmap, color-scale is normalized to the max trial averaged firing rate achieved in the case of the respective walk neuron activation simulation. Arrow thickness indicates synaptic strength (5-400 synapses). Red: excitatory, blue: inhibitory connections. Asterisks indicate neurons (oDN1, BDN2) selected for further analysis in Figure 3. **(e)** Connectome based prediction of halting neurons. Dark red filled circles indicate empirical verified halt neurons and light red filled circles indicate predicted halt neurons. Dark green filled circles indicate empirical walk neurons and light green filled circles indicate predicted walk neurons. Arrow thickness indicates synaptic strength (5-400 synapses). Red: excitatory, blue: inhibitory connections. **(f)** EM segmentation of predicted halt neurons in the brain. **(g)** Input zone brain region for predicted halt neurons.

**Supplementary Video 1.** Transient activation of different halt neurons (BB, FG, BRK) using CsChrimson in tethered flies with annotated leg joints (top) and corresponding 3D reconstruction of leg kinematics (bottom), related to Extended Data Fig 3. Red circle on top left indicates light ON (625 nm LED). Video is being played at 50% speed.

**Supplementary Video 2.** Transient activation of P9 alone or co-activated with different halt neurons (FG, BB, FG+BB) using CsChrimson in free-walking flies, related to Figure 2. Red dot on top left indicates light ON (630 nm LED).

**Supplementary Video 3.** DNs downstream of P9, color-coded according to their firing rate (top right) after *in silico* stimulation of P9 alone or co-activated with FG or BB, related to Figure 3. Asterisk show neuron of interest (oDN1).

**Supplementary Video 4.** DNs downstream of BPN, color-coded according to their firing rate (top right) after *in silico* stimulation of BPN alone or co-activated with FG or BB, related to Figure 3. Asterisk show neuron of interest (BDN2).

**Supplementary Video 5.** Calcium activity recording (2P) of BDN2 axons in the ventral nerve cord of a BDN2-Gal4>Act88F:Rpr; UAS-GC6f;tdTomato fly during walking, related to Figure 3. White arrowheads indicate BRK axons (middle), red moving bar on plot (right) shows translational velocity synchronized with fly video (left) and imaging recording.

**Supplementary Video 6.** Transient activation of MDN or BDN2 alone (left), and MDN or BDN2 co-activated with BRK (right) using CsChrimson in decapitated, tethered flies with annotated leg joints and corresponding 3D reconstruction of leg kinematics, related to Figure 4. Red circle on top left indicates light ON (625sss nm LED). Video is being played at 50% speed.

**Supplementary Video 7.** Calcium activity recording (1P from 0-15s, 2P from 16-30s) of femoral muscles in the front leg of MHC-LexA>LexAOP-GCaMP6f; BRK-Gal4>UAS-CsChrimson flies with their Fe-Ti joint forcibly kept (glued) in extended (left) or flexed position (right), related to Figure 4. Red dot indicates light ON (655nm LED), filled arrowheads show muscle activity on BRK activation, empty arrowheads indicate inactive muscles. Video is being played at 50% speed.

**Supplementary Video 8.** Calcium activity recording (2P) of femoral muscles in the front leg of MHC-LexA>LexAOP-GCaMP6f; BRK-Gal4>UAS-CsChrimson flies with their Fe-Ti joint forcibly kept (glued) in extended (left) or flexed position (right), related to Figure 4. Red dot indicates light ON (655nm LED), filled arrowheads show muscle activity on BRK activation, empty arrowheads show inactive muscles/ inhibition of pre-stimulation muscle activity. Video is being played at 50% speed.

**Supplementary Video 9.** Calcium activity recording (2P) of BRK axons in the ventral nerve cord of BRK-Gal4>Act88F:Rpr; UAS-GC6f tdTomato flies during front leg rubbing (Fly1, left) and hind leg rubbing (Fly2, right), related to Figure 6. Colored squares indicate onset of rubbing events, white arrowheads indicate segment specific BRK activity. Video is being played at 20% speed.

**Supplementary Video 10.** Transient inhibition in grooming intact flies (from 0-7s), and decapitated flies (from 8-22s), for control and BRK>GtACR1 flies, related to Figure 6. Green dot at top left indicates light ON (540nm LED).

**Supplementary Table 1.** Excel file including all abbreviated genotypes used in this study, their full experimental genotypes and exact sample sizes, grouped by figure numbers and panels.

**Supplementary Table 2.** Excel file including all strains used in this study, as well as their full genotypes, abbreviations and sources/identifiers.

**Supplementary Table 3.** Excel file including all FlyWire/FANC/MANC identifiers for left- and right-hemisphere neurons considered in this study, as well as their community labels, cell types and names given in this study.

